# SiPhyNetwork: An R package for Simulating Phylogenetic Networks

**DOI:** 10.1101/2022.10.26.513953

**Authors:** Joshua A. Justison, Claudia Solis-Lemus, Tracy A. Heath

## Abstract

1. Gene flow is increasingly recognized as an important macroevolutionary process. The many mechanisms that contribute to gene flow (*e*.*g*., introgression, hybridization, lateral gene transfer) uniquely affect the diversification of dynamics of species, making it important to be able to account for these idiosyncrasies when constructing phylogenetic models. Existing phylogenetic-network simulators for macroevolution are limited in the ways they model gene flow.
2. We present SiPhyNetwork, an R package for simulating phylogenetic networks under a birth-death-hybridization process.
3. Our package unifies the existing birth-death-hybridization models while also extending the toolkit for modeling gene flow. This tool can create patterns of reticulation such as hybridization, lateral gene transfer, and introgression.
4. Specifically, we model different reticulate events by allowing events to either add, remove, or keep constant the number of lineages. Additionally, we allow reticulation events to be trait-dependent, creating the ability to model the expanse of isolating mechanisms that prevent gene flow. This tool makes it possible for researchers to model many of the complex biological factors associated with gene flow in a phylogenetic context.

## 1 Introduction

Interspecific gene flow—the movement of genetic material across species boundaries—is observed throughout the tree of life (Mallet et al., 2016). Interspecific gene flow (gene flow hereafter) processes such as the transmission of genes without vertical inheritence to parents (lateral-gene transfer), interbreeding between species (hybridization), and backcrossing between hybrids and their parental lineages (introgression), can operate across wide ranges of both genetic and taxonomic scales. These dynamic processes can facilitate the sharing of genetic material as small as single genes or as large as whole chromosomes or genomes. Further, exchanges happen not only between closely related populations and species complexes, but also between organisms from different kingdoms of life. These events can have a profound effect on species and lineages at micro- and macroevolutionary scales, playing a significant role in mimicry complexes (Smith and Kronforst, 2013; Enciso-Romero et al., 2017), invasion ecology (Rhymer and Simberloff, 1996; Viard et al., 2020), insecticide resistance (Norris et al., 2015), and adaptive radiations (Grant and Grant, 2019; Meier et al., 2017, 2019; Moest et al., 2020). Regardless of the mode of gene flow and the many ways in which it may affect reticulate species, it is clear that it is an important factor in shaping macroevolutionary patterns (Stebbins, 1959; Bock, 2010; Taylor and Larson, 2019).

With the increased accessibility and availability of genomic data for lineages across the tree of life, an increasing number of studies have found the signatures of historical reticulation and gene flow (Taylor and Larson, 2019). These studies use a wide range of methods for detecting gene flow and describing the processes responsible for generating patterns of reticulation. The available approaches for untangling past gene flow vary widely in how these events are characterized and the scope and scale to which they can be applied—some methods estimate the presence of gene flow, while others seek to estimate when gene flow occurred and which parts of the genome have reticulate histories (see Payseur and Rieseberg, 2016). Recent advances allow researchers to directly estimate gene flow through phylogenetic network inference, with approaches ranging from parsimony and distance-based methods to model-based likelihood and Bayesian methods (Elworth et al., 2019).

Although the field has made significant progress in the development of phylogenetic-network inference methods, the tools for simulating networks under relevant macroevolutionary processes remain limited. Simulated data are vital for validating the accuracy and examining the performance of statistical methods. Moreover, simulation tools can also be useful in empirical studies for hypothesis testing and evaluating model adequacy. Due to the limited availability of phylogenetic network simulators, network-based simulation studies often rely on using bifurcating trees with randomly added reticulate edges (*e*.*g*., Bastide et al., 2018; Hejase et al., 2018), simulating sequence data from empirically derived networks (*e*.*g*., Wen et al., 2016), or by using an arbitrary fixed phylogenetic network (*e*.*g*., Solís-Lemus and Ané, 2016; Wen and Nakhleh, 2018). While these approaches to creating datasets with known attributes are useful for testing specific scenarios and highlighting core features of methods, these networks are not generated by biologically relevant or stochastic model–based processes, limiting the range of conditions that can be explored and rendering the conclusions less generalizable.

Birth-death processes are often used to describe macroevolutionary patterns (Kendall, 1948; Nee, 2006) and, consequently, are also commonly used in phylogenetic simulators (*e*.*g*., Stadler, 2011; Höhna, 2013; Höhna et al., 2015; Hagen and Stadler, 2018). To further model and simulate under important mechanisms of biological systems, many extensions of the birth-death process have been developed. These extensions include density-dependent diversification (Rabosky and Lovette, 2008), time-dependent rates (Stadler, 2011; Höhna, 2013), lineage age-dependent rates (Hagen and Stadler, 2018), and fossilization (Barido-Sottani et al., 2019). Extensions of the birth-death process even simulate phylogenetic networks by allowing hybridization events (Morin and Moret, 2006; Woodhams et al., 2016; Zhang et al., 2018). Although some macroevolutionary simulators can incorporate gene flow, each simulator makes different assumptions about how reticulation events affect the phylogeny (Figure 1; Table 1).

**Table 1:**
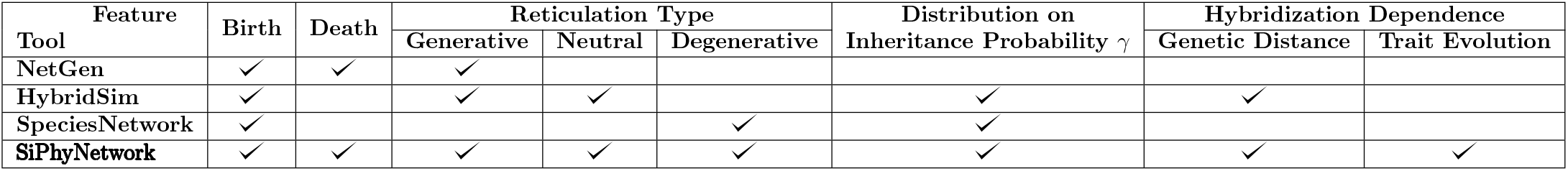
Simulation model features present in existing tools employing the birth-death-hybridization process. Reticulation type is denoted as generative (denoted as m-type in Janssen and Liu, 2021), neutral (n-type), and degenerative (y-type), referring to the net gain, stasis, or reduction in the number of lineages from an event, respectively (see Figure 1). A distribution on the inheritance probability *γ* enables users to specify an arbitrary distribution to determine the inheritance proportions of the parental lineages at hybridization. Hybridization dependence allows the success of hybridization to rely on either genetic distance or on a trait that evolves on the phylogenetic network.

| Feature | Birth | Death | Reticulation Type |  |  | Distribution on | Hybridization Dependence |  |
| --- | --- | --- | --- | --- | --- | --- | --- | --- |
| Tool | | | Generative | Neutral | Degenerative | Inheritance Probability $\gamma$ | Genetic Distance | Trait Evolution |
| NetGen | ✓ | ✓ | ✓ |  |  |  |  |  |
| HybridSim | ✓ |  | ✓ | ✓ |  | ✓ | ✓ |  |
| SpeciesNetwork | ✓ |  |  |  | ✓ | ✓ |  |  |
| **SiPhyNetwork** | ✓ | ✓ | ✓ | ✓ | ✓ | ✓ | ✓ | ✓ |

**Figure 1:**
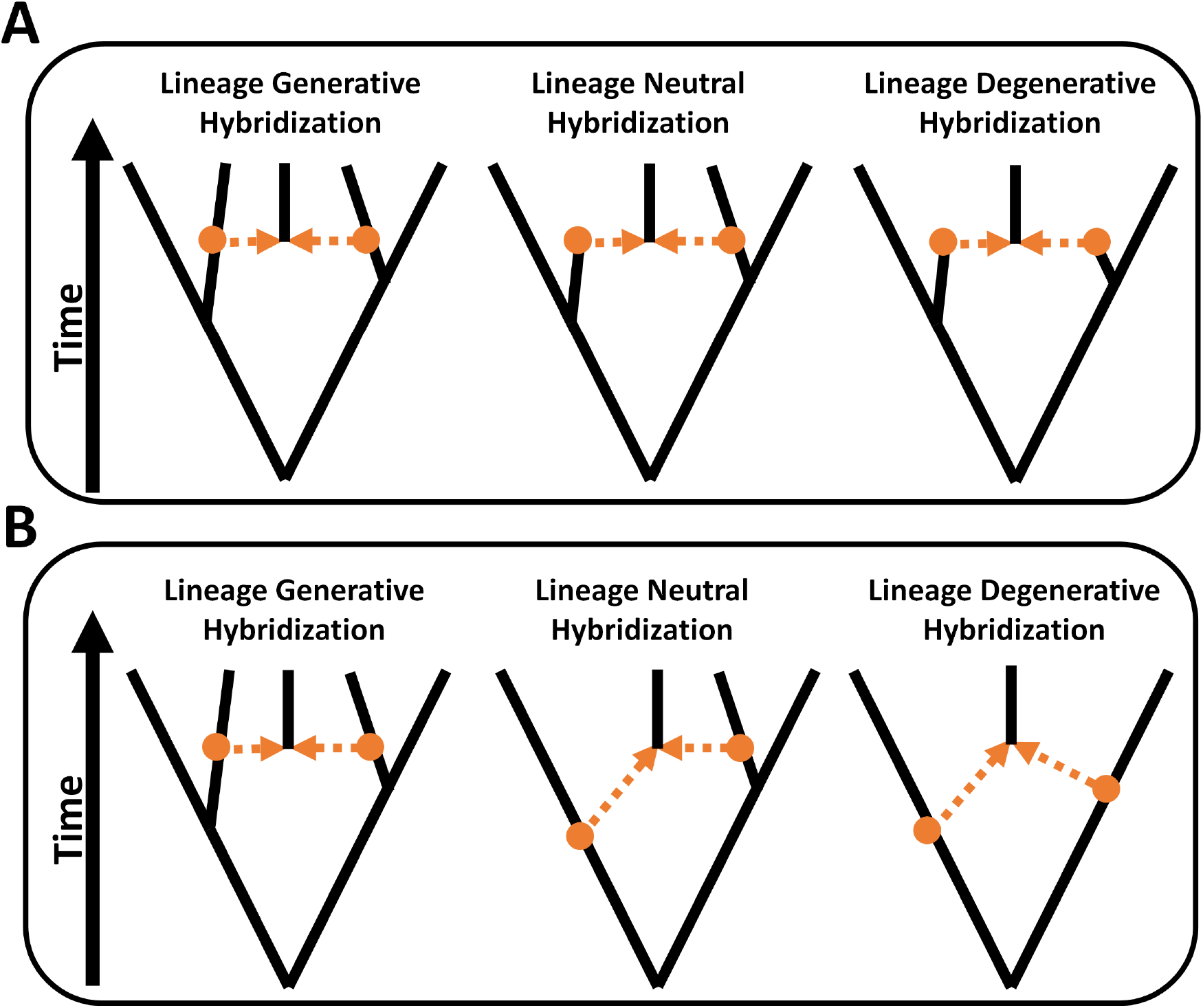
Macroevolutionary patterns of gene flow. Orange circles denote the parental nodes that lead to the reticulate node, while dashed orange arrows indicate the two parental lineages that contribute to gene flow. Lineage generative hybridization (m-type) occurs when a reticulation event results in a gain of one lineage. Lineage neutral hybridization (n-type) results in the a net-zero change in the number of lineages. Lineage degenerative hybridization (y-type) reduces the number of lineages by one. Reticulation events on phylogenetic networks are typically drawn using one of two conventions: A) having all parental nodes lateral to the hybrid node, or B) fusing any in-degree 1 out-degree 1 nodes, potentially placing parent nodes at different times than the hybrid node. Both depictions, however, have the same topological relationships.

Simulation tools generate phylogenetic networks using a variety of models such as the coalescent (Arenas and Posada, 2007; Kelleher et al., 2016; McKenzie and Eaton, 2020), genome evolution models (Mallo et al., 2016; Davín et al., 2020), or generate networks based on certain phylogenetic characteristics (Janssen and Liu, 2021). While these processes are often useful for modeling population genetic and genomic processes, birth-death processes are invoked to describe lineage diversification on a macroevolutionary scale. HybridSim (Woodhams et al., 2016), NetGen (Morin and Moret, 2006), and the SpeciesNetwork package in BEAST2 (Zhang et al., 2018) all simulate networks under variants of the birth-death-hybridization process, which models how lineages speciate, go extinct, and hybridize as a stochastic process. However, it is important to recognize the different model assumptions about the diversification process and different conceptualizations of hybridization found in each of the available simulators (Table 1). For example, extinction has a considerable effect on shaping biodiversity yet only NetGen explicitly models extinction events while simulating phylogenetic networks. Although, other simulators do not model the loss of species, they have additional flexibility in how they handle gene flow events by having different types of gene flow and allowing parental lineages to differentially contribute to the hybrid species. The disparity of event types and how certain events are modeled between simulators, in turn, make the evaluation of patterns in networks challenging, since the distribution of generated networks is dependent on the inputs and assumptions of available simulation software. As such, an important addition to the present toolkit is a simulator capable of generating phylogenetic networks from a unified framework, allowing for more direct comparisons between the effects of these assumptions on generated networks.

The evolution of morphological characters and other phenotypic traits can additionally lead to reproductive isolation under a variety of pre-zygotic and post-zygotic mechanisms, and consequently, act as barriers to to hybridization (Grant, 1981; Soltis and Soltis, 2009; Abbott et al., 2013). For example, studies have described traits imposing reproductive isolation in lice where body size differences cause mechanical isolation (Villa et al., 2019), butterflies selectively mating with individuals that match their mimetic coloration (Dincă et al., 2013), or many plant species where chromosomal rearrangements create post-pollination barriers (Baack et al., 2015). Even viable hybrids may become excluded on macroevolutionary scales due to low fitness of subsequent generations. This phenomenon, termed hybrid breakdown (Grant, 1981; Soltis and Soltis, 2009), can occur if the hybrid has an intermediate trait value and is unable to find a niche different from the parental lineages, resulting in lower fitness.

Although HybridSim is capable of modulating the amount of hybridization in relation to the genetic distance between lineages, no tools currently account for the effect that certain morphological characters can have on the ability for lineages to hybridize.

Here we present the R package SiPhyNetwork that enables phylogenetic network simulation under a range of biological scenarios, and extends the currently available tools by:

1. modeling the different ways that reticulation can affect lineages during gene flow events,
2. allowing simulations to have trait-dependent hybridization,
3. adapting many common tree simulation features and utility functions for networks (*e*.*g*., incomplete lineage sampling, complete versus extant only phylogeny, sampling under the generalized sampling approach of Hartmann et al., 2010),
4. unifying many unique model features from other model simulators in one package (*i*.*e*., asymmetric inheritance and genetic distance dependent hybridization as seen in HybridSim), and
5. providing functions for manipulating and classifying phylogenetic networks.

Although reticulation on a macroevolutionary scale is typically described with hybridization events, SiPhyNetwork does not make any specific assumptions about the mechanism of gene flow. Consequently, SiPhyNetwork can model other reticulate processes like introgression or lateral gene transfer. Overall, we sought to provide a framework for simulating evolutionary histories under a multitude of reticulate mechanisms found across the Tree of Life. We believe that this work will enable researchers to test a wide array of hypotheses about reticulate macroevolution.

## 2 Model Components and Implementation

SiPhyNetwork is an R (R Core Team, 2022) package that simulates phylogenetic networks under a birth-death-hybridization process. Our implementation is a generalization of the constant-rate birth-death-hybridization process that allows hybridization to either add a lineage (lineage generative), remove a lineage (lineage degenerative), or keep the number of lineages constant (lineage neutral)—unifying the models of Woodhams et al. (2016) and Zhang et al. (2018) in a single framework. These types of hybridization events are also denoted m-type, y-type, and n-type reticulations respectively (Janssen and Liu, 2021). We additionally allow trait-dependent hybridization and genetic distance–dependent hybridization. SiPhyNetwork is available on the Comprehensive R Archive Network (CRAN). Alternatively, the source code and installation instructions for the development version can be accessed at https://github.com/jjustison/SiPhyNetwork.

Network simulation using SiPhyNetwork relies on three core functions: `sim.bdh.age()`, `sim.bdh.taxa.ssa()`, and `sim.bdh.taxa.gsa()`, collectively referred to as the `sim.bdh()` functions. All three functions use the same simulation algorithm but have different stopping conditions (discussed below). With the exception of arguments setting the stopping condition, each simulation function takes the same set of arguments to specify the model (Table 2). These functions generate `evonet` objects from the ape software package (Paradis and Schliep, 2019), which are themselves extensions of the `phylo` objects used for phylogenetic trees. These objects can then be stored in the extended Newick format (Cardona et al., 2008) so they can be used for downstream macroevolutionary analyses (*e*.*g*., Solís-Lemus et al., 2017; Bastide et al., 2018) or visualization (*e*.*g*., Vaughan, 2017; Schliep et al., 2021). In the following sections, we demonstrate how the arguments are used for each component of the birth-death-hybridization model. Further details about the R implementation and examples can be found in the “SiPhyNetwork Introduction” vignette that is released with the package and included in the supplementary materials.

**Table 2:** Arguments of simulation functions in SiPhyNetwork

| **Parameter** | **Description** |
| --- | --- |
| `age, n, m` | Stopping conditions |
| `numsim` | The number of simulation replicates |
| `lambda` | Speciation rate ($\lambda$) |
| `mu` | Extinction rate ($\mu$) |
| `nu` | Hybridization rate ($\nu$) |
| `hybprops` | Hybridization-type proportions |
| `hyb.inher.fxn` | A function that determines inheritance probabilities |
| `hyb.rate.fxn` | A function that relates genetic distance to hybridization success |
| `trait.model` | A list containing a model for trait evolution and hybridization-dependence rules for the traits |
| `frac` | The sampling fraction of extant species |
| `stochsampling` | A logical that if **TRUE**, then each extant taxon is sampled with probability **frac**. If **FALSE**, then a constant proportion of **frac** taxa are sampled, rounded to the nearest whole taxon |
| `twolineages` | A logical that if **TRUE**, starts the process with two lineages that share a common ancestor, else starts the process with one lineage |
| `complete` | A logical to return the complete or reconstructed network |

### 2.1 Diversification Process

We model the branching process of the phylogeny with speciation, extinction, and hybridization events. The lineage diversification process has exponentially distributed waiting times for the events, with constant-rate parameters *λ* for speciation, *μ* for extinction, and *ν* for hybridization. Since hybridization requires two lineages, we consider the rate of hybridization (*ν*) on each species pair, whereas speciation (*λ*) and extinction (*μ*) are rates on each lineage. For a phylogeny with *N* taxa, a rate on each species pair means that hybridization events occur at an effective rate of 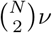 The overall waiting time until the next event of the birth-death-hybridization process is exponentially distributed with rate

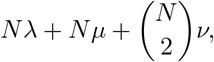

where the probability for each event is weighted by its effective rate for *N* taxa (*i*.*e*., *Nλ* for speciation, *Nμ* for extinction, and 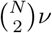 for hybridization). In SiPhyNetwork users specify a value for each rate (`lambda`, `mu`, and `nu`) by providing values in the `sim.bdh()` functions.

For each hybridization event, we denote the genetic contributions of the two parental lineages—also called inheritance probabilities—as *γ* and 1 *− γ*. Inheritance probabilities are drawn from a user-defined distribution that draws values from 0 to 1, allowing for asymmetric inheritance, where one parent contributes more genetic material, and broad flexibility to match prior beliefs about gene flow. For example, supplying a Beta(10, 10) distribution can model hybrid speciation where inheritance probabilities are largely equal, while a Beta(0.1, 0.1) would reflect introgression where one parental lineage often contributes a larger proportion of genetic material (Figure 2). In SiPhyNetwork, users supply a function for the `hyb.inher.fxn` argument in the `sim.bdh()` functions to draw inheritance probabilities at each hybridization event. There are several helper functions in SiPhyNetwork that create functions for inheritance probabilities, as shown in the example below.

**Figure 2:**
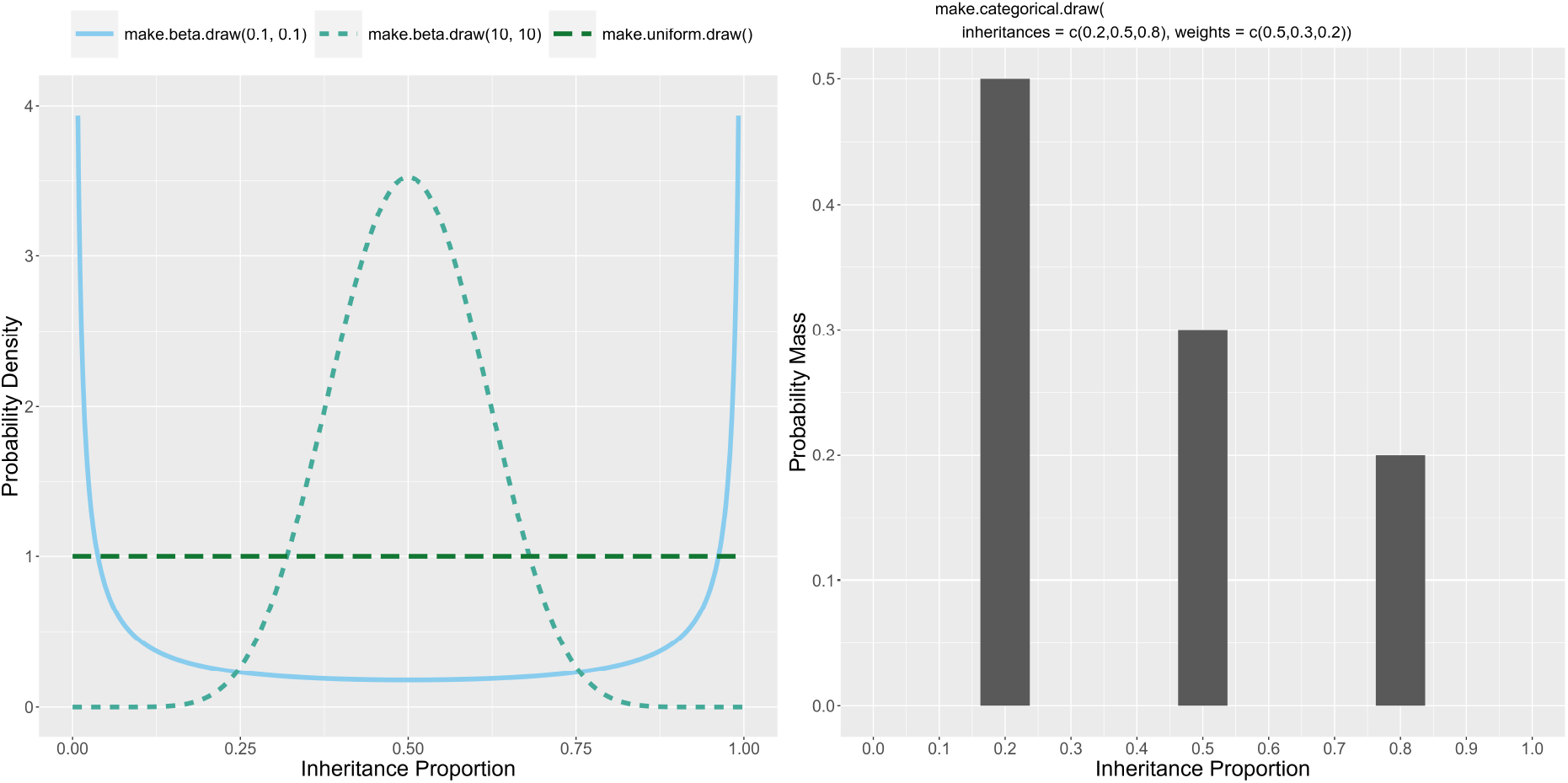
Examples of sampling distributions for inheritance proportions *γ* and the corresponding functions in R to make the distributions. Inheritance proportions are drawn from the sampling distribution at each hybridization event. Users can specify any distribution or set of values as long as the output is between 0 and 1. The distributions on the left depict continuous sampling distributions on [0, 1] while the distribution on the right only draws values 0.2, 0.5, or 0.8

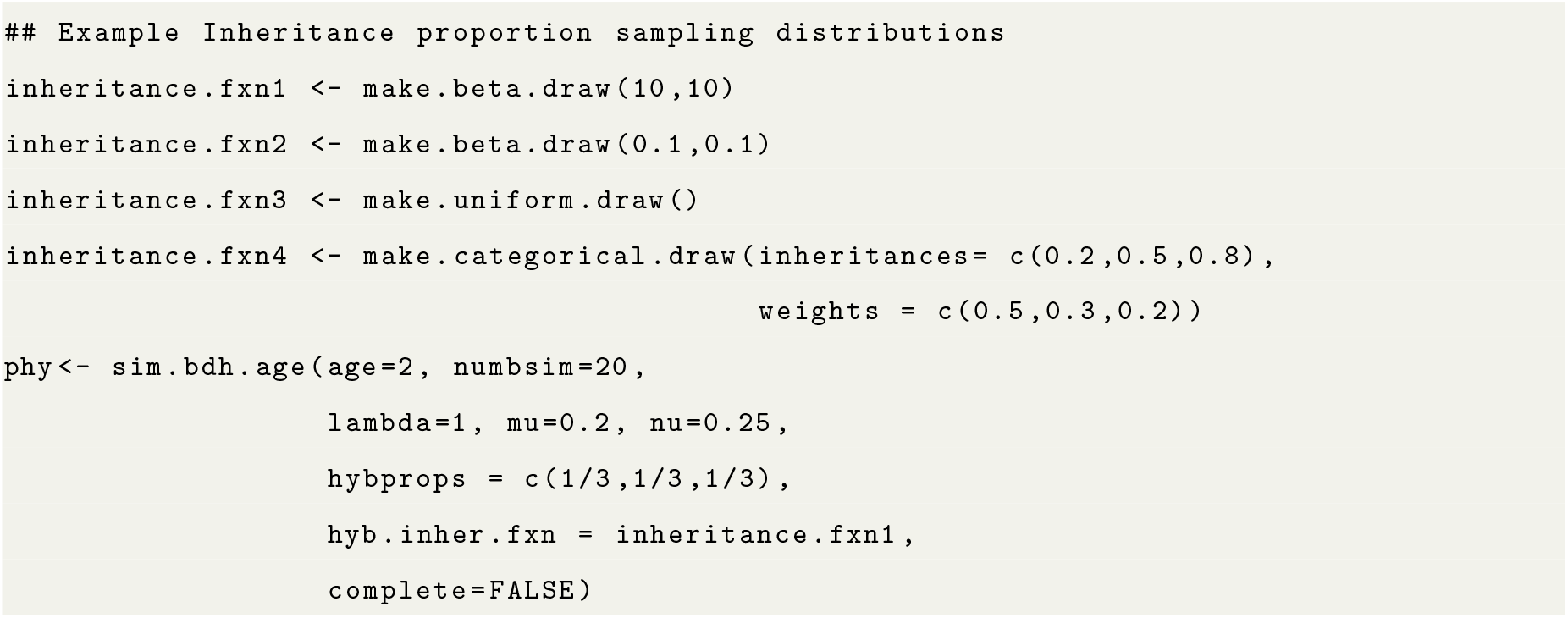

Additionally, users can create their own functions to sample inheritance probabilities when hybridization events occur. The supplied function should take no arguments and return a number between 0 and 1.

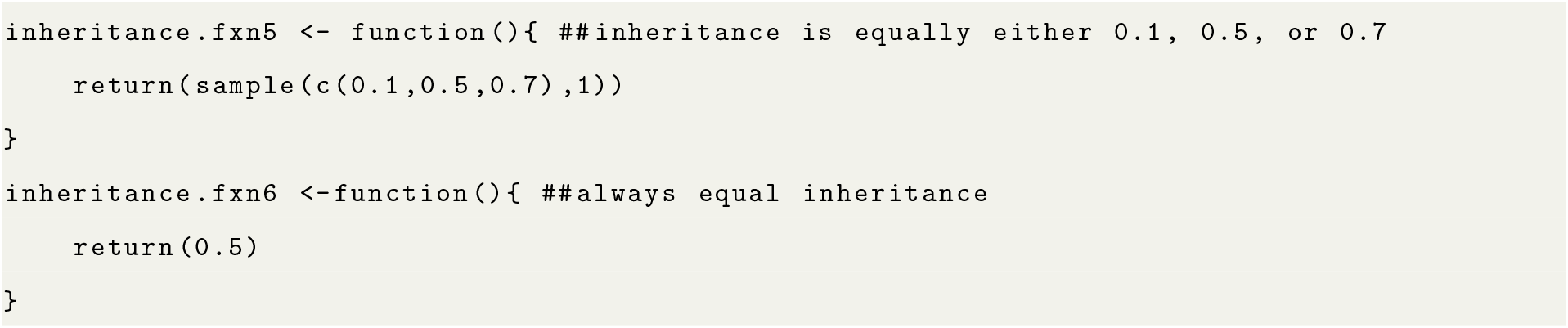

### 2.2 Hybridization Type and the Effect on the Number of Lineages

SiPhyNetwork has a versatile system for simulating many modes of gene flow by allowing for reticulation events with different macroevolutionary patterns (see Figure 1). We denote each type of event by the net change in the number of lineages as a result of the hybridization: lineage generative when gaining a lineage, lineage neutral when maintaining the same number of lineages, and lineage degenerative for when a lineage is lost. Each type of hybridization imposes different time constraints on the parent nodes (orange circles in Figure 1) that lead to the reticulate node. Both parental nodes co-occur with the reticulation for lineage generative events, only one parent occurs at the same time as the reticulate event for lineage-neutral events, and there are no time co-occurrence constraints for lineage-degenerative events. When a hybridization event occurs, it is either lineage generative, lineage neutral, or lineage degenerative, with probabilities *ρ*_+_, *ρ*_0_, and *ρ*_*−*_, respectively.

Users have the ability to specify the probabilities for each macroevolutionary pattern, giving the flexibility to model various gene flow types and microevolutionary mechanisms. Although we do not make mechanistic assumptions about how gene flow occurs, certain processes may be better at describing a given reticulate pattern. For example, modeling hybrid speciation with lineage generative hybridization would be appropriate due to the creation of a new hybrid lineage on the phylogeny. However, lineage-neutral and lineage-degenerative hybridization may also be valid models for hybrid speciation in the cases where genetic swamping occurs (Todesco et al., 2016) or in the presence of ghost lineages (Ottenburghs, 2020; Tricou et al., 2022). Additionally, lineage-neutral hybridization could be used to model cases of introgression or lateral-gene transfer, in which gene flow occurs but no new lineages are produced.

Existing phylogenetic network simulators allow different subsets of these reticulation patterns (Table 1). Netgen solely considers lineage-generative hybridization (Morin and Moret, 2006), HybridSim considers lineagegenerative and lineage-neutral hybridization, with the latter being termed ‘introgression’ (Woodhams et al., 2016), and the SpeciesNetwork package considers solely lineage-degenerative hybridization (Zhang et al., 2018).

Hybrid-type probabilities are modeled in SiPhyNetwork by providing a vector of probabilities for each pattern of hybridization. Each of the elements in the vector corresponds to the probability that hybridization is generative, neutral, or degenerative, respectively. The vector is given in the `hybprops` argument of the `sim.bdh` functions. Below we show examples of different specifications for the hybrid-type proportions.

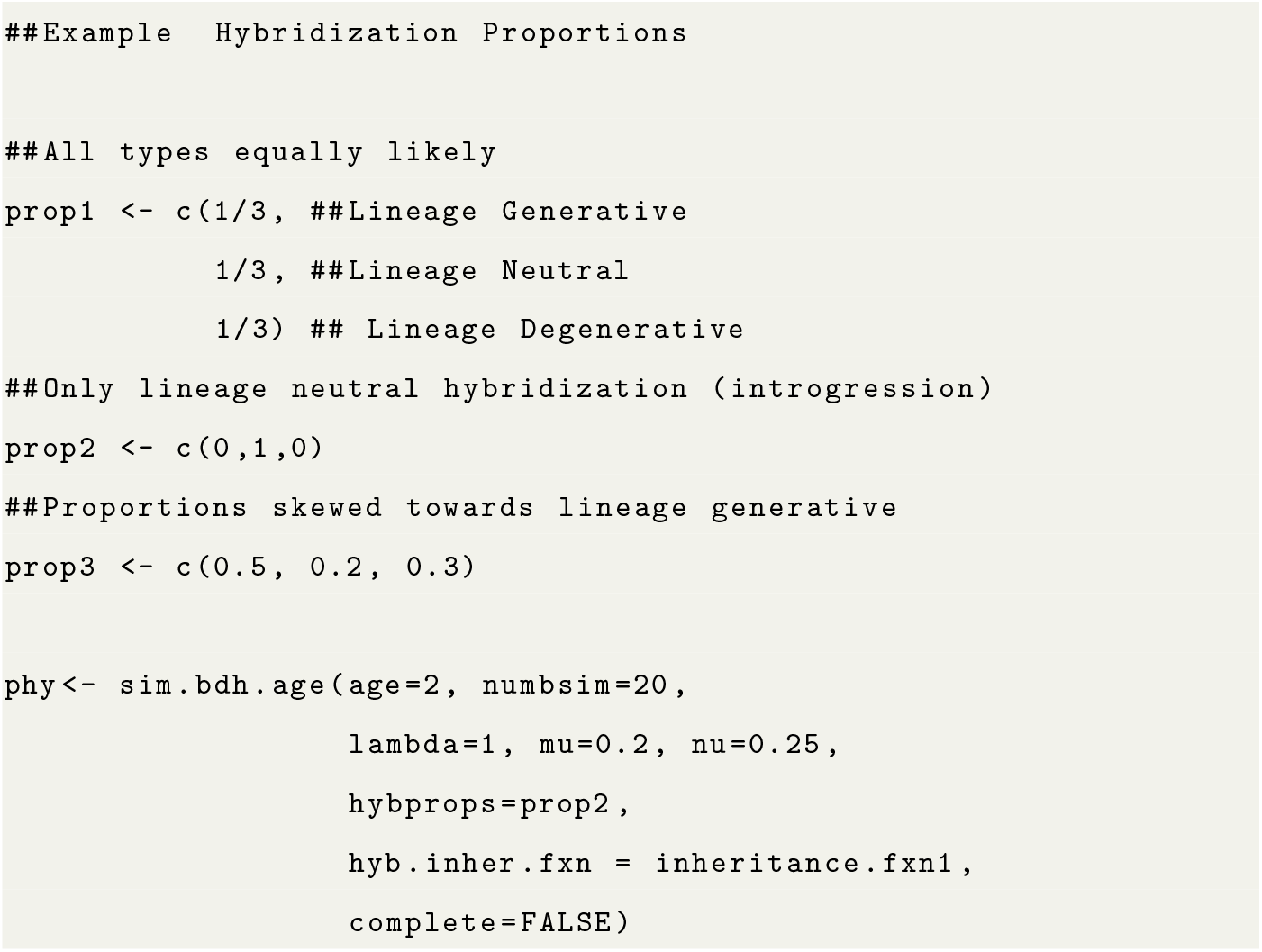

### 2.3 Hybridization Success Dependent on Genetic Distance

Gene flow occurs more frequently between closely related lineages than it does for distantly related lineages (Gourbiere and Mallet, 2010; Abbott et al., 2013). We model hybridization success as a function of genetic distance between lineages in SiPhyNetwork using the approach of Woodhams et al. (2016). Hybridization events effectively become a non-stationary Poisson process with respect genetic distance. The hybridization rate changes as a function of the genetic distance *d*_*ij*_ between taxa *i* and *j* at a given time:

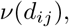

where the relationship between hybridization and genetic distance is user-specified. However, in practice we use the thinning of a Poisson process to break this into two steps: (1) hybridization events between a given species pair are proposed as part of a Poisson process with rate *ν* and (2) proposed hybridization events are then successful with a probability that is proportional to the genetic distance between the species pair. Successful events are added to the phylogeny while unsuccessful hybridization attempts are not. A genetic distance matrix is maintained throughout the simulation and is updated at each event during the forward-in-time simulation to accurately reflect genetic distances at any given point in time. The genetic distance between two taxa *i* and *j* is the total length of edges that are not shared on the path from each taxon to the root (or a weighted summation of each path if the taxon in question has hybrid ancestry). Formally, we assume a strict molecular clock where the genetic distance at a given time is denoted as:

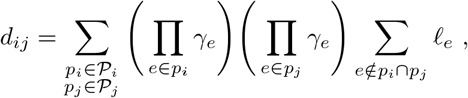

where *P*_*i*_ denotes the set of paths from the taxon *i* to the root, *γ*_*e*_ is the inheritance probability of the associated edge *e*, and *𝓁*_*e*_ is the edge length of *e*. This formulation is identical to the covariance computation of Bastide et al. (2018) with the exception that we take the sum of edge lengths that are not shared across paths, instead of taking the edges that are shared.

In SiPhyNetwork genetic-distance dependence is modeled by providing a genetic-distance function for the `hyb.rate.fxn` argument. Users have the flexibility to define any arbitrary function that relates hybridization success to genetic distance. This function takes the genetic distance as an argument and should return a number that represents the probability of hybridization success, as shown in the example below.

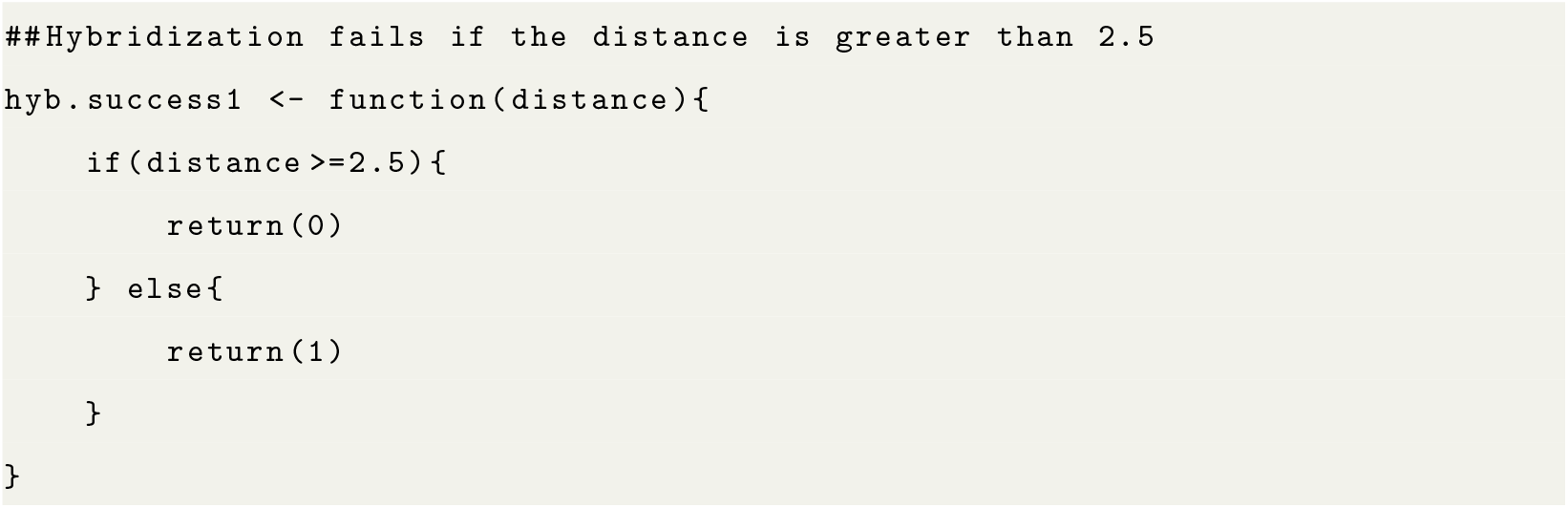

Additionally, we have implemented the same decreasing functions as Woodhams et al. (2016), *i*.*e*., linear decay, exponential decay, snowballing decay, and polynomial decay:

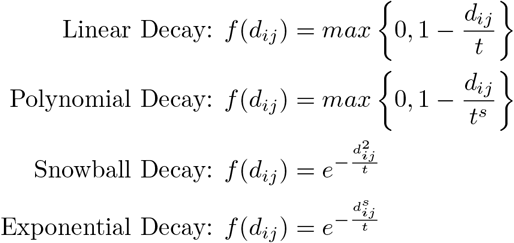

Here, *s* and *t* are values set by the user that are used to affect the shape and rate of decay for each function. We have several helper functions to create these genetic-distance dependence functions, as shown below

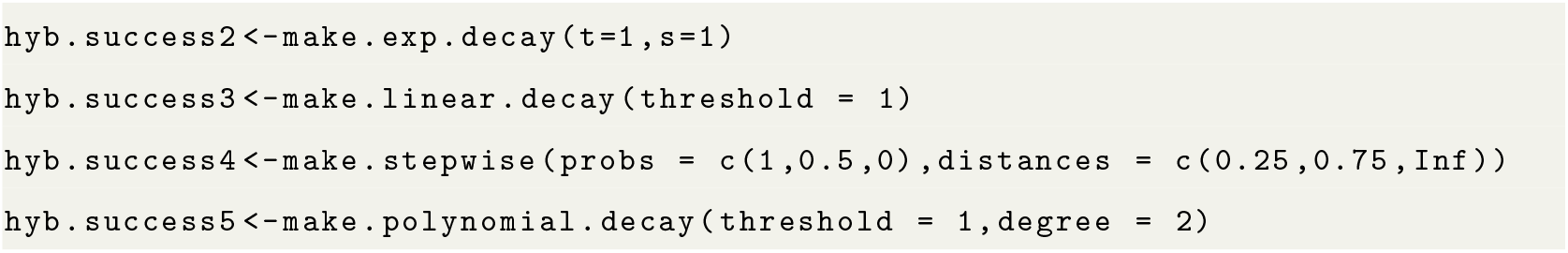

### 2.4 Trait-Dependent Hybridization

In SiPhyNetwork, we implemented a general framework for modeling the complex interplay between successful hybridization and trait evolution. Our trait-dependent hybridization model has three components: a trait-evolution model, a model for trait inheritance in hybrid lineages, and rules that describe how hybridization success depends on trait values (Figure 3). The model of trait evolution specifies how continuous or discrete trait values change over time and has the flexibility to implement a number of trait evolution models (*e*.*g*., Brownian motion (Felsenstein, 1985), Ornstein–Uhlenbeck (Lande, 1980), Mk (Pagel, 1994; Lewis, 2001), threshold (Felsenstein, 2005)). The model for trait inheritance specifies how the trait is inherited both at speciation events and at hybridization events. The last component permits the user to define how trait values interact to determine whether hybridization occurs. In nature, both discrete and continuous traits are known to affect rates of hybridization, thus SiPhyNetwork allows either type of phenotypic trait to affect the hybridization potential of different lineages. Likewise, in some systems, traits that are more similar may enhance the likelihood of hybridization (Dincă et al., 2013; Pereira et al., 2014), while in others, opposite trait values create novelty for hybrids to succeed (Vereecken et al., 2010).

**Figure 3:**
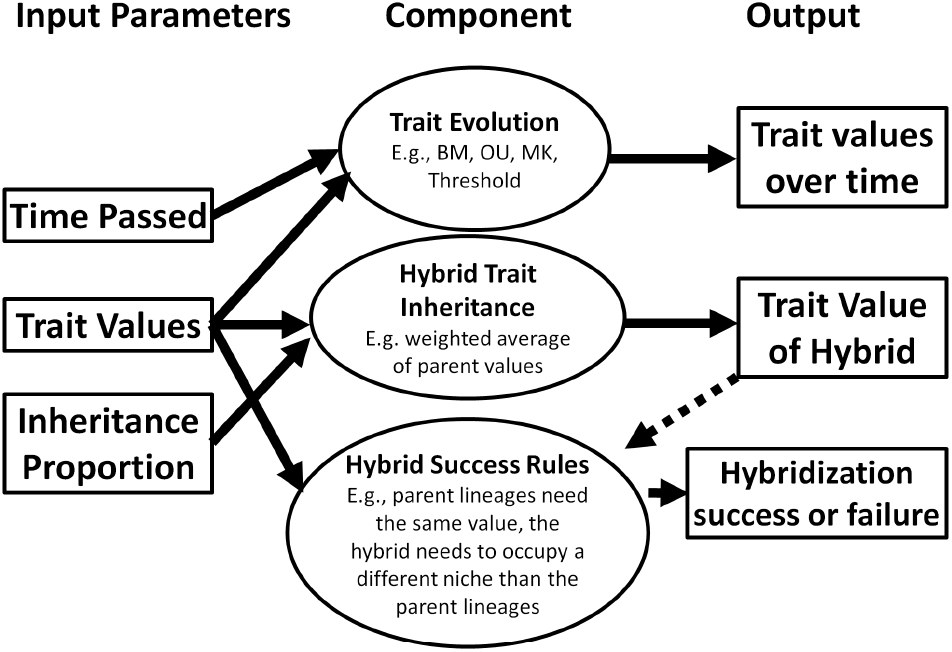
A schematic diagram of the SiPhyNetwork model for trait-dependent hybridization. Each column depicts the model inputs, model components, and outputs of each model, respectively. Arrows leading into each model component denote the needed inputs and the arrows leading out of each model component depict the outputs. The trait value of the hybrid—indicated with a dashed arrow—is a special output that can also be used an optional input for determining hybrid success. The model for trait-dependent hybridization has three sub-components: the trait evolution model, hybrid trait inheritance, and rules for hybridization success. The first component describes how trait values change over time for each lineage. The second models how the hybrid inherits its trait value from its parents. The last component enforces rules that determine whether the hybridization event is successful and occurs on the phylogeny or fails and does not occur. Further, if desired, the last component uses the trait value of the hybrid from the second component as an input for determining hybridization success.

Each component is created with user-defined functions to determine how they operate during simulation. This framework offers a great degree of flexibility for modeling biologically realistic trait-dependent hybridization scenarios. For example, one can generate networks of hybridizing lineages modulated by ploidy evolution with both allo- and auto-polylploidization; characters can evolve continuously such that a hybrid can only persist if it avoids hybrid breakdown by occupying a trait space different than that of its parental lineages (Soltis and Soltis, 2009); or hybridization could become less successful as the traits of two lineages become increasingly dissimilar. Thus, SiPhyNetwork is flexible in modeling trait-dependent hybridization by allowing the biologist to tailor the mode to their particular system.

We model trait-dependent hybridization by supplying the optional argument `trait model` to the `sim.bdh()` functions. The `trait model` argument is a list that specifies each component of the trait-dependent hybridization model (Figure 3). The trait model is a named list with the following elements: `initial states` a value for the initial state of the trait, `hyb.event.fxn` a function to determine how the trait is inherited on the hybrid lineage, `hyb.compatability.fxn` a function to determine whether hybridization can occur based on the trait values, `time.fxn` a function that determines how the traits change over time, and `spec.fxn` a function that determines how the trait is inherited at speciation events. In the example below we implement a model for ploidy evolution that considers autopolyploidy and allopolyploid hybridization. We restrict allopolyploidy events to only occur between lineages with the same ploidy.

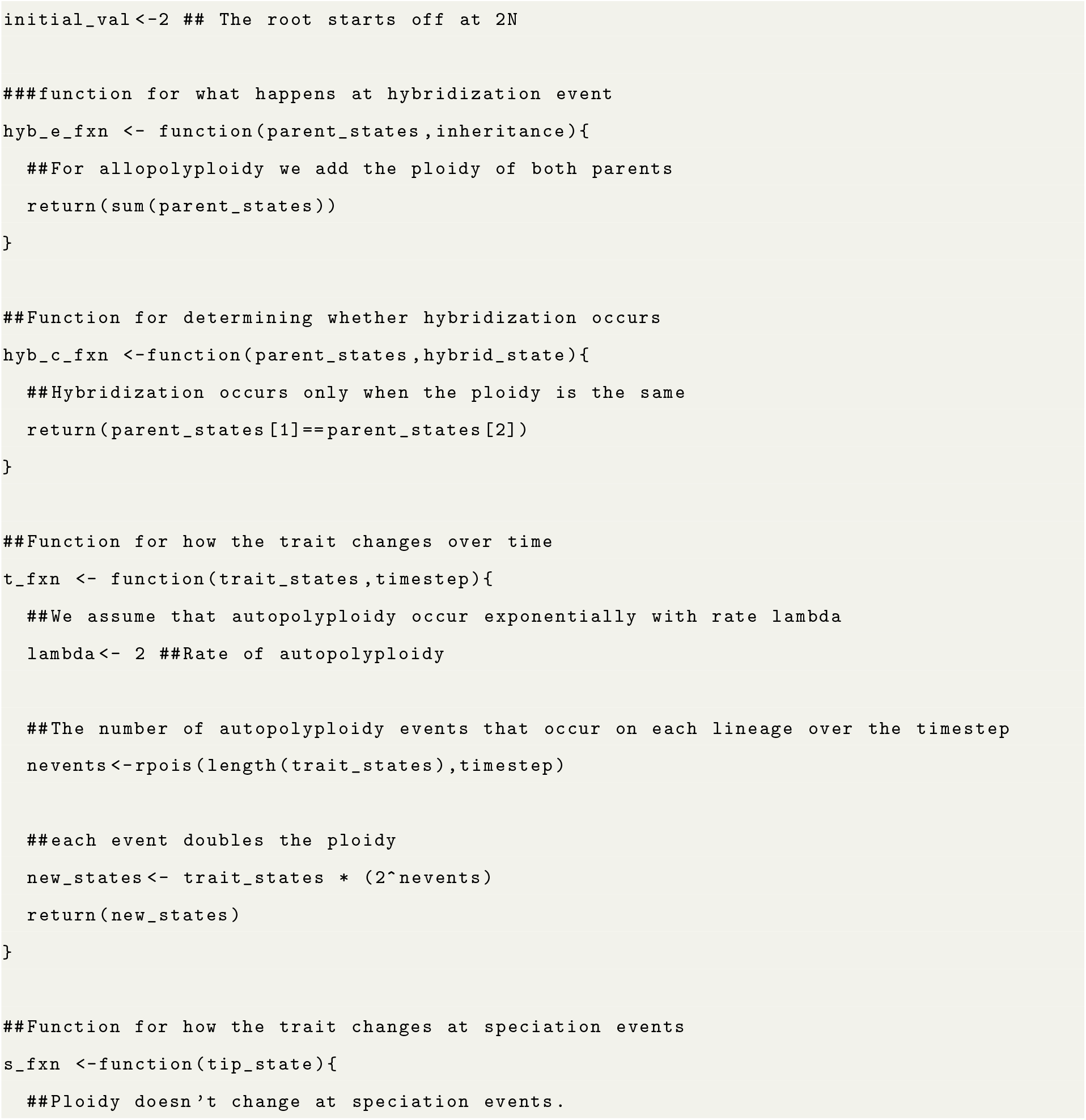
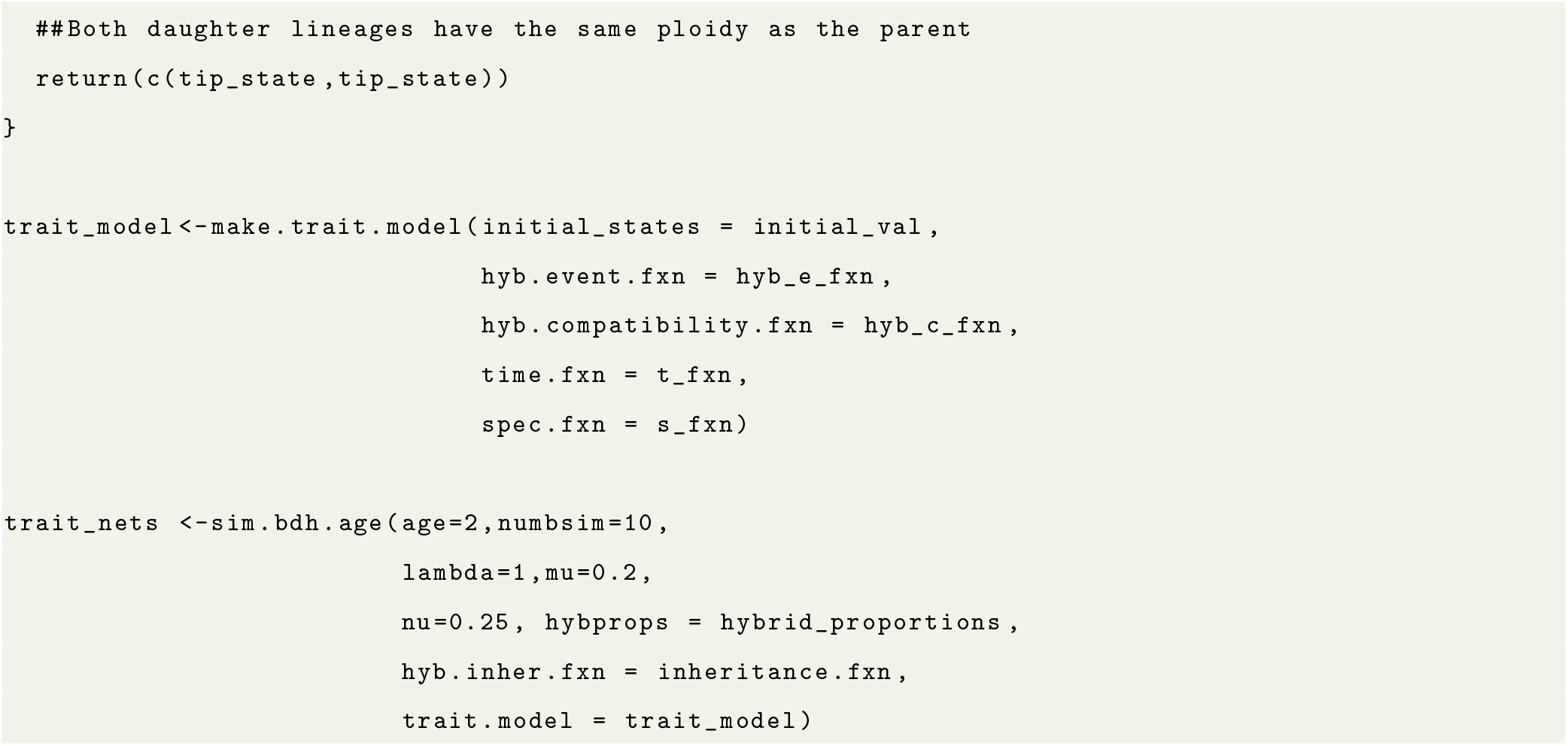

More information about model capability and specific implementations can be found in the “Introduction” vignette (Supplemental files).

### 2.5 Extant-Only and Incomplete Sampling

Typically, phylogenetic networks do not include all extant taxa or they lack fossil specimens that can provide information about extinct lineages. We can model incomplete lineage sampling by pruning away unsampled lineages (Figure 4), leaving what is often referred to as the reconstructed or sampled phylogenetic network (Gernhard, 2008; Stadler, 2009). Indeed, it is necessary to account for incomplete sampling as it affects expected branch-length distributions (Nee et al., 1994; Stadler, 2008). Producing an extant-only phylogeny is a common feature of phylogenetic tree simulators, but not available in current phylogenetic network simulators. We extend these features to phylogenetic networks, both as a core part of phylogenetic network simulation and as a *post-hoc* operation on phylogenetic networks with utility functions. SiPhyNetwork models these processes in the `sim.bhd()` functions by setting `complete=FALSE` to eliminate all extinct taxa and setting `frac` to less than 1 for incomplete sampling of extant taxa (Figure 4). If `frac` is less than one, then that proportion of extant taxa will be sampled. Further, if the argument `stochsampling=TRUE`, then each extant taxon will be sampled with probability `frac`. If `stochsampling=FALSE`, then `frac` proportion of taxa will be sampled from the phylogeny, rounded to the nearest whole number.

**Figure 4:**
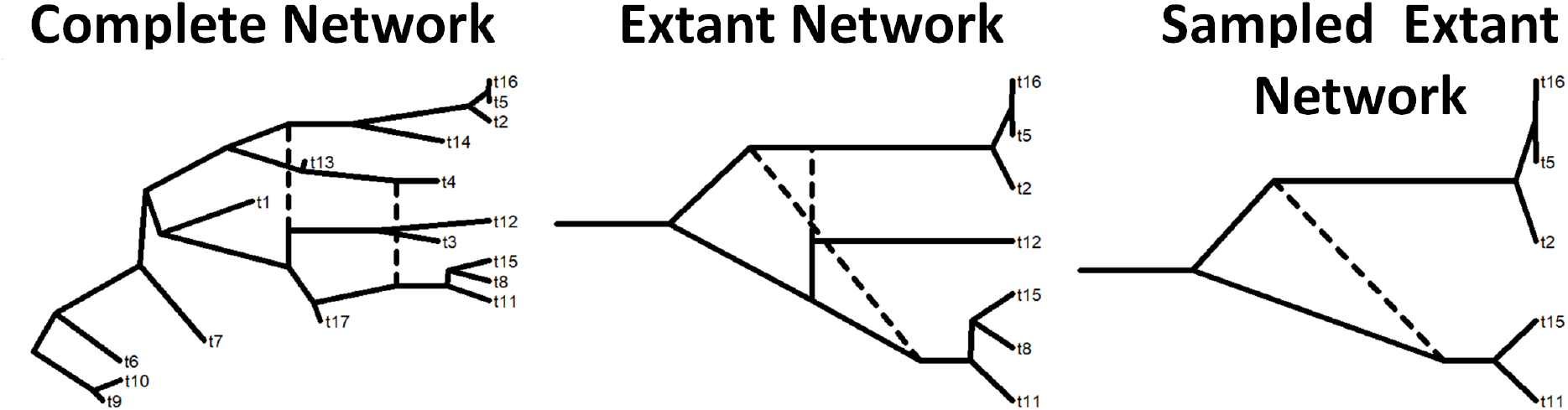
Lineage sampling on a phylogenetic network from SiPhyNetwork. Here we show the same phylogenetic network under three lineage-sampling procedures. The complete network represents the entire simulated history and includes all extant and extinct lineages. The extant network shows only the history of the extant lineages after pruning all extinct lineage. Interestingly, although the complete network has a lineage neutral hybridization at MRCA(t15, t8, t11), the same hybridization appears as a lineage degenerative hybridization in the extant phylogenetic network. The sampled extant network shows the reticulate history of a subset of the extant taxa after pruning all extinct lineages and the extant taxa *t*8 and *t*12 pruned. Without the hybrid lineage *t*12 in the sampled extant network, the lineage-generative hybridization is no longer observable.

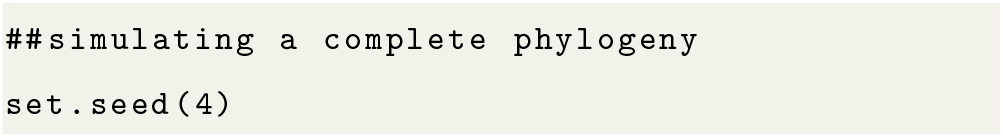
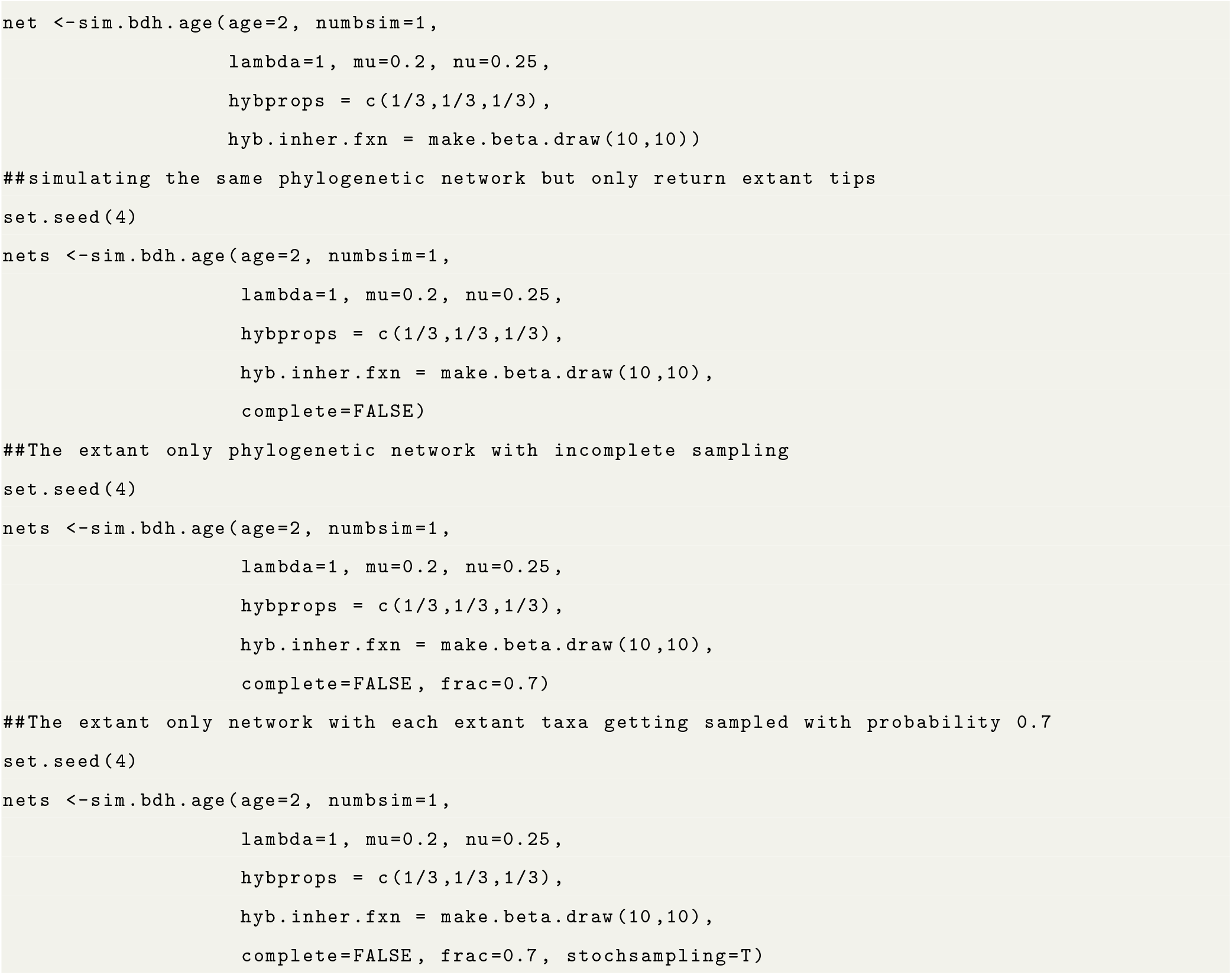

### 2.6 Sampling Strategies

Users have two options for defining a simulation’s stopping condition, where the simulation ends once (1) a specified time or (2) a specified number of taxa is reached. Traditionally, phylogenies simulated to a number of taxa allow the process to continue until first reaching *N* taxa, known as the simple sampling approach (SSA). However, this approach does not correctly sample to a number of taxa while assuming a uniform prior on tree ages, and doing so is a not a trivial task (Hartmann et al., 2010; Stadler, 2011).

We extend the generalized sampling approach (GSA) for generating birth-death trees that was introduced by Hartmann et al. (2010) to phylogenetic network simulation, which correctly samples networks with a specified number of taxa under the birth-death-hybridization process. Briefly, if *N* taxa are desired under the GSA, the simulation process will continue until reaching *M* taxa, then phylogenies are uniformly sampled from periods with *N* taxa. A sufficiently large value of *M* (*i*.*e*., *M >> N*) should be chosen such that the probability of the process returning to *≤ N* taxa is small. The function `sim.bdh.age()` simulates the process from the origin until a specified age, while `sim.bdh.taxa.ssa()` and `sim.bdh.taxa.gsa()` are used to simulate to a specified number of taxa under the SSA and GSA approaches, respectfully. Birth-death simulations can routinely go extinct before reaching a stopping condition or never reach a desired number of species in a tractable amount of time under certain parameterizations (*e*.*g*., *μ > λ*). Similarly, for the birth-death-hybridization process, the specific combinations of *λ, μ, ν* values will affect the probability that a simulation reaches its stopping condition.

## 3 Discussion

SiPhyNetwork brings much of the currently available phylogenetic-network simulation functionality into a single R package, while also extending existing models by considering various macroevolutionary patterns of reticulation and allowing for trait-dependent hybridization. The different types of gene flow (lineage generative, neutral, and degenerative) have received some attention, although primarily through the context of the timing of hybridization events (Hibbins and Hahn, 2019; Flouri et al., 2020). The constraints produced by each gene-flow type pose an interesting yet challenging problem for inference (Hibbins and Hahn, 2022). Since each type of hybridization necessitates a different number of speciation events to explain the same number of lineages, over-attributing a specific type of hybridization likely would lead to bias in diversification-rate estimates. Additionally, both sampling only extant taxa (Nee et al., 1994) and incomplete sampling (Stadler, 2008, 2009), are known change our expectations about the birth-death process and resulting distributions of bifurcating trees. However, it is not well characterized how these processes change our expectations of the birth-death-hybridization process. In fact, incomplete sampling and the presence of ghost lineages can make it particularly difficult to infer the correct reticulate pattern (Ottenburghs, 2020; Tricou et al., 2022). Additionally, failure to sample parental lineages can remove the node co-occurrence constraint in the extant-only phylogeny, making it appear as another type of hybridization when compared to the complete phylogeny (Figure 4).

The birth-death-hybridization process and other biological extensions may be able to explain certain macroevolutionary patterns. First, ancient gene flow is frequently found in empirical studies (Meier et al., 2017; Pavón-Vázquez et al., 2021; Zhang et al., 2021). Yet, under the simple birth-death-hybridization process, ancient gene flow should be rare compared to contemporary gene flow due to there being fewer species and the effective hybridization rate scaling more quickly per taxon than speciation or extinction. Hybridization dependent on genetic distance or certain characteristics may make the effective hybridization rate scale more slowly to explain the pattern of ancient gene flow. Additionally, high lineage-degenerative hybridization may have the ability to explain the slowdown in lineage accumulation, often attributed to density-dependent diversification (Rabosky and Lovette, 2008) or time-dependent diversification (Hagen and Stadler, 2018). In this case, lineage accumulation slows down because the rate of lineage-degenerative hybridization scales more quickly with the number of taxa than the rate of speciation. Eventually the lineage-degenerative hybridization would reduce the net-diversification rate to zero until reaching and revolving around some steady state of taxa.

SiPhyNetwork is a tool that facilitates our understanding of patterns of reticulate diversification. Further, the birth-death-hybridization process has many unique properties that we have only begun to explore and this work allows further characterization by sampling from the distribution of phylogenetic networks under this macroevolutionary process. Moreover, SiPhyNetwork provides a framework to test and validate inference methods under a stochastic and biologically informed model that accounts for many gene flow processes.

## Supporting information

Introduction Vignette

## 4 Authors’ Contributions

Joshua A. Justison, Claudia Solis-Lemus, and Tracy A. Heath designed the models. Joshua A. Justison implemented the methods. Joshua A. Justison and Claudia Solis-Lemus tested the methods. Joshua A. Justison, Claudia Solis-Lemus, and Tracy A. Heath wrote the manuscript.

## 5 Acknowledgements

We would like to thank folks from the Heath lab and Solis-Lemus lab for helpful comments and discussion while developing this tool. We would additionally like to thank Cecile Ane and the many other users that submitted bug reports and provided feedback on the software in its early states. Lastly, we would like to thank the editors, Florian C. Boucher, and three other anonymous reviewers for helpful comments and review. This work was partially funded by the National Science Foundation [DEB-2144367 to CSL].

## 7 Data Accessibility Statement

SiPhyNetwork GPL v3 open-source license held by Joshua Justison, Claudia Solis-Lemus and, Tracy A. Heath. SiPhyNetwork is tested on current and future versions of R. All code is avaliable on Github: https://github.com/jjustison/SiPhyNetwork and on Zenodo (Justison, 2023).

