## Supplementary material for "SiPhyNetwork: An R package for Simulating Phylogenetic Networks": Introduction Vignette

Josh Justison

### Overview

The SiPhyNetwork R package is a tool for simulating phylogenetic networks under birth-death-hybridization based models. It provides functions for creating complex hybridization models, functions for simulating phylogenies under these models, as well as utility functions for characterizing the types of phylogenetic networks, visualizing networks, and saving phylogenetic networks in the Extended Newick format. This allows users to simulate phylogenetic networks under a suite of more biologically realistic models. The SiPhyNetwork package is released under the GPL-3 license and can be downloaded from CRAN or the source code is avaliable on GitHub

### Introduction

Birth-death processes are commonly used to describe species diversification *(Nee 2006)*. There exist several phylogenetic simulation tools that generate trees under the birth-death process *(See Stadler 2011)* as well as extensions for density-dependent diversification , time-dependent rates *(Höhna 2013; Höhna, May, and Moore 2015)*, lineage age dependent rates *(Hagen and Stadler 2018)*, and fossilization *(Barido-Sottani et al. 2019)*.

Gene flow and hybridization is found throughout the tree of life and has had a nontrivial role in shaping diversity *(Mallet, Besansky, and Hahn 2016)*. Recently, an extension of the birth-death process that includes hybridization was developed as a prior for Bayesian phylogenetic inference *(Zhang et al. 2018)*. This model assumes the waiting time between hybridizations is exponentially distributed with rate parameter $\binom{ntaxa}{2}\nu$, where $\nu$ is the hybridization rate. A few tools exist that simulate phylogenetic networks under variants of this model *(Morin and Moret 2006; Woodhams, Lockhart, and Holland 2016; Davín et al. 2020)*, however they all have different conceptualizations on how hybridization events affect the number of lineages on the phylogeny. As process-based phylogenetic inference methods receive increased attention, it becomes imperative that we can simulate biologically relevant phylogenetic network for the testing of these methods and for creating hypotheses about empirical systems.

### The SiPhyNetwork Package

The SiPhyNetwork R package allows simulation of phylogenetic networks under a birth-death-hybridization process. This package extends the capabilities of other existing birth-death-hybridization based simulators by allowing:

- Simulating to a fixed time or fixed number of species
- Incomplete sampling sampling
- Sampling of extinct taxa

- Hybridization events to be either a lineage generating, a lineage neutral, or a lineage degenerative process.

- Successful hybridization events to be an arbitrary function of genetic distance
- Successful hybridization events to be dependent on a trait that evolves along the network
- Functions for visualizing, writing, and reading phylogenetic networks in R

Throughout this vignette we will cover how to use these features in SiPhyNetwork.

### Simulating Networks

There are three core functions for simulating phylogenetic networks: `sim.bdh.age`, `sim.bdh.taxa.ssa`, and `sim.bdh.taxa.gsa` (collectively referred to as `sim.bdh` style functions). These functions all use the same birth-death-hybridization models but differ in their stopping conditions.

- The `sim.bdh.age` function simulates until a specified `age`.
- The `sim.bdh.taxa.ssa` function uses the Simple Sampling Approach (SSA) to simulate to a specified number of taxa `n`, that is, the simulation will stop once the phylogeny has `n` tips.
- The `sim.bdh.taxa.gsa` function uses the General Sampling Approach (GSA) to simulate to a specified number of taxa `n`. Briefly, the GSA simulates `m` taxa under the SSA and samples the phylogeny from time periods where `n` taxa are present.

Both the SSA and GSA are described with greater detail in *Hartmann, Wong, and Stadler (2010)* and *Stadler (2011)*.

#### Example 1: Simulating Networks Under A Simple Hybridization Model

The `sim.bdh` style functions have many optional arguments with default values. For this first set of simulations we won't adjust any optional arguments, we'll be playing with these arguments in later sections. However, it is important to know the assumptions we are making with our model by using default parameters. Specifically, we assume:

- `frac = 1` and `stochsampling = FALSE` assumes that all extant taxa are sampled in the phylogeny.
- `twolineages=FALSE` means that we start with a one lineage rather than two lineages that share a common ancestor
- `complete = TRUE` leaves extinct species on the phylogeny.
- `hyb.rate.fxn = NULL` assumes that successful hybridization events are not a function of the genetic distance between taxa.
- `trait.model = NULL` assumes that successful hybridization events do not depend on a trait value between taxa.

The arguments we will be using are:

- `age`, `m`, and `n`: These parameters are used for the stopping condition of the simulations.
- `numbsim`: The number of simulations performed
- `lambda`: The speciation rate
- `mu` The extinction rate
- `nu` The hybridization rate
- `hybprops`: A vector of length three that denotes the proportion of hybridizations that are lineage generative, lineage degenerative, and lineage neutral.
- `hyb.inher.fxn`: This is a function used for determining the inheritance proportions.

Now we can try running some simulations

```
library(ape)
library(SiPhyNetwork)
set.seed(82589933) ##set the seed for reproducibility.
##This is the exponent of the largest known Mersenne prime number

##First we need a function that describes how inheritance probabilities are drawn
inheritance.fxn <- make.beta.draw(10,10)
##We can see that this function makes draws from a beta(10,10) distribution
inheritance.fxn()
#> [1] 0.4893861
inheritance.fxn()
#> [1] 0.4677096

##We also want to set the proportion of each type of hybrid event
hybrid_proportions <-c(0.5,  ##Lineage Generative
                       0.25, ##Lineage Degenerative
                       0.25) ##Lineage Neutral

##We can simulate to 7 extant tips with the SSA
ssa_nets<-sim.bdh.taxa.ssa(n=7,numbsim=20,
                    lambda=1,mu=0.2,
                    nu=0.20, hybprops = hybrid_proportions,
                    hyb.inher.fxn = inheritance.fxn)
ssa_net<-ssa_nets[[20]] ##The sim.bdh functions return a list of length numbsim. We get the 20th simula
print(ssa_net)
#>
#>     Evolutionary network with 2 reticulations
#>
#>                --- Base tree ---
#> Phylogenetic tree with 8 tips and 12 internal nodes.
#>
#> Tip labels:
#>   t2, t4, t7, t6, t9, t8, ...
#>
#> Rooted; includes branch lengths.

##We can also simulate 7 extant taxa with the GSA.
##We choose m=30 because it becomes very unlikely that at 30 tips we will ever return to 7
gsa_nets<-sim.bdh.taxa.gsa(m=30,n=7,numbsim=20,
                    lambda=1,mu=0.6,
                    nu=0.3, hybprops = hybrid_proportions,
                    hyb.inher.fxn = inheritance.fxn)
gsa_net<-gsa_nets[[19]]

##Simulate a network up to age 2
age_nets <-sim.bdh.age(age=2,numbsim=20,
                    lambda=1,mu=0.2,
                    nu=0.25, hybprops = hybrid_proportions,
                    hyb.inher.fxn = inheritance.fxn)
age_net<-age_nets[[8]]
```

There are a few things worth noting here. Firstly, if we looked at `age_nets` we would notice that some of the elements are `0`, the `sim.bdh` style functions return `0` if the phylogeny goes extinct before reaching the stopping condition.

Secondly, we can see that each phylogeny is an `evonet` object, however, they have an additional attribute `inheritance`. `inheritance` contains a vector of inheritance probabilities. The $i^{th}$ element in `inheritance` corresponds to the inheritance probability of the hybrid edge denoted in the $i^{th}$ row of the `reticulation` attribute.

```
age_net$inheritance ##This corresponds to the edges found in reticulation
#> [1] 0.3674308 0.3716242 0.4939105 0.5539802
age_net$reticulation
#>      from to
#> [1,]    9 10
#> [2,]   15 18
#> [3,]   17 21
#> [4,]   22 20
```

Further, we can use the `gsa.network()` function to sample phylogenies under the GSA. This allows us to properly sample $n$ taxa from any phylogenetic model, regardless if the phylogenetic networks were generated from the `SiPhyNetwork` package or not.

```
##We can simulate to 30 extant tips under the SSA.
##In this case the 30 acts as the m parameter of the GSA
ssa_nets<-sim.bdh.taxa.ssa(n=30,numbsim=10,
                    lambda=1,mu=0.2,
                    nu=0.20, hybprops = hybrid_proportions,
                    hyb.inher.fxn = inheritance.fxn)

my_net<-ssa_nets[[1]]
my_net<-network.gsa(net=my_net,ntaxa=5,complete=T,frac=1,stochsampling=F)
```

Lastly, Most of the arguments in the `sim.bdh` type functions take either a numeric or boolean, however, `hyb.inher.fxn`, `hyb.rate.fxn` take functions as arguments while `trait.model` takes a list of functions as an argument. These take functions as arguments so the user can define functions as they please to model the specific biology of interest. Some utility functions exist to aid in creating functions that fit the specific purpose of these arguments.

For example, when simulating above, we used `make.beta.draw(10,10)` to create a function that makes draws from a beta distribution with shape parameters 10 and 10. We might chose a beta function like this if we believed that inheritance proportions are generally equal but have some variation. Alternatively, `make.uniform.draw()`, and `make.categorical.draw()` are other utility functions for creating appropriate functions for the `hyb.inher.fxn` argument. In fact, any function will work as long as that function requires no arguments itself and returns values on the range $[0, 1]$.

We will explore how to appropriately make arguments for `hyb.rate.fxn` and `trait.model` later in this vignette.

#### Types of Hybridization

We often think of hybridization as a species forming event, a new hybrid species gets created. Although a new hybrid species gets created, the net number of lineages doesn't need to strictly increase. We should also consider cases where one or both of the parental lineages go extinct as a result of the hybridization. The hybrid species could either outcompete one or both of the parental lineages and displace them or the hybrid species could continually backcross with the parental lineages until one homogenous lineage remains, a process known as genetic assimilation. If one parental lineage goes extinct we have zero change in the number of lineages while if both parental lineages go extinct we have a net loss of one lineage even with the creation of the hybrid species. We call these different outcomes lineage generative, lineage neutral, and lineage degenerative hybridization.

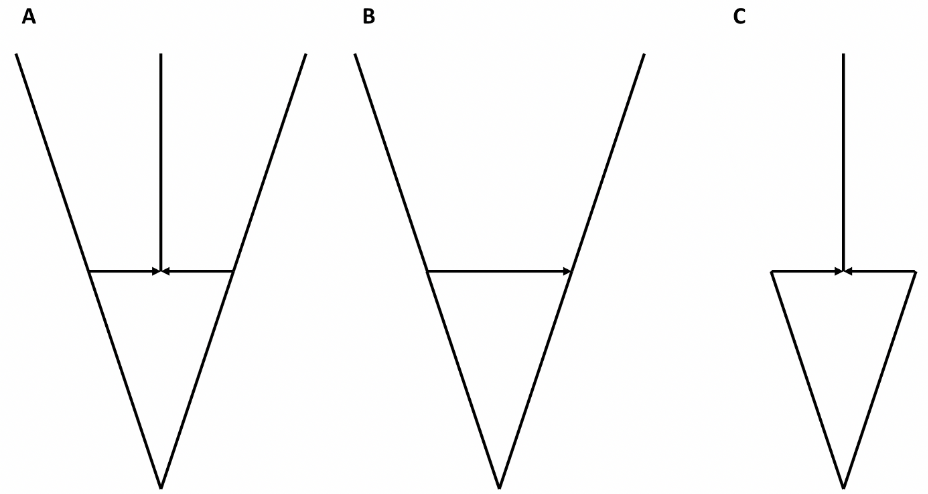


*Different Notions of Hybridization. Subfigures depict* ***A)*** *lineage generating hybridization (speciating hybridization),* ***B)*** *lineage neutral hybridization, and* ***C)*** *lineage degenerative hybridization*

**Example 2 Types of Hybridization in SiPhyNetwork**

Modeling the different types of hybridization in SiPhyNetwork is relatively straightforward with use of the `hybprops` argument in the `sim.bdh` functions. `hybprops` is a vector of length three that denotes the relative probability that a hybridization event is lineage generative, lineage degenerative, or lineage neutral respectively.

```
##Equal chance of all three types
hybprops1 <-c(1/3, ##Lineage Generative
              1/3, ##Lineage Degenerative
              1/3) ##Lineage Neutral

##Skewed chance of all three types
hybprops2 <-c(0.5, ##Lineage Generative
              0.2, ##Lineage Degenerative
              0.3) ##Lineage Neutral

##Only Lineage Generative Hybridization occurs
hybprops3 <-c(1, ##Lineage Generative
              0, ##Lineage Degenerative
              0) ##Lineage Neutral


##simulate where all 3 are equally likely
age_nets <-sim.bdh.age(age=2,numbsim=20,
                    lambda=1,mu=0.2,
                    nu=0.25, hybprops = hybprops1,
                    hyb.inher.fxn = inheritance.fxn)
```

### Characterizing And Using Phylogenetic Networks

After simulating phylogenetic networks it can be helpful to identify certain characteristics of the networks and eventually save those networks to file so they can be used in other pipelines.

#### Example 3 Plotting Networks

One of the first ways of understanding the type of phylogenetic networks we generated would be to plot them

```
plot(ssa_net,main="SSA Network")
plot(gsa_net,main="GSA Network")
#> Warning in min(x): no non-missing arguments to min; returning Inf
#> Warning in max(x): no non-missing arguments to max; returning -Inf
plot(age_net,main="Age Network")
```

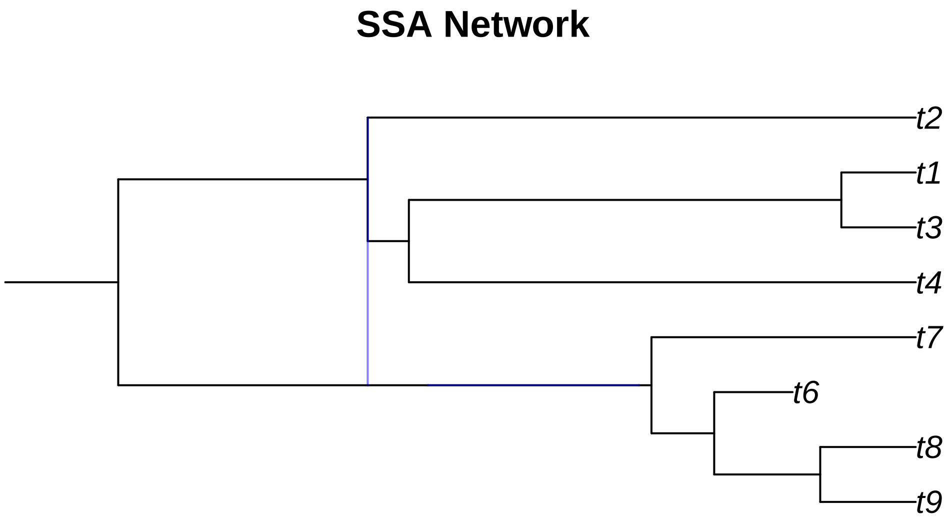

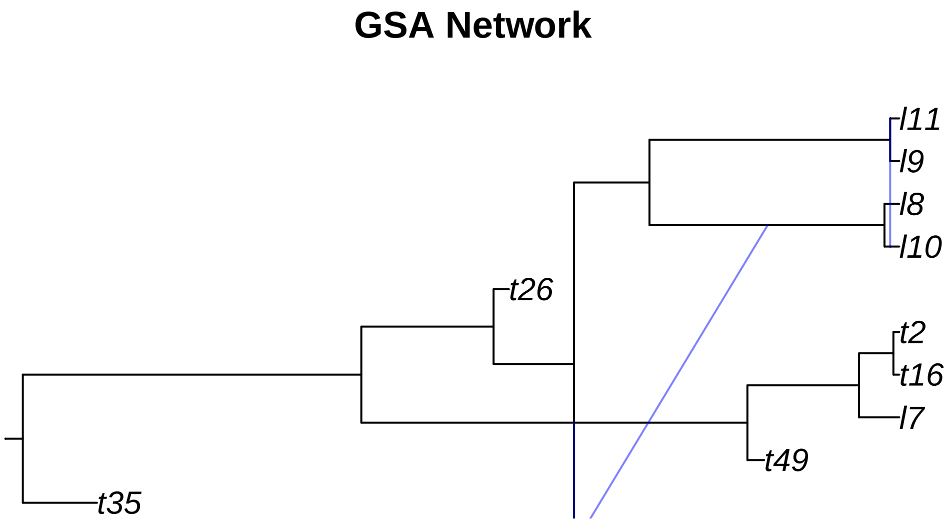


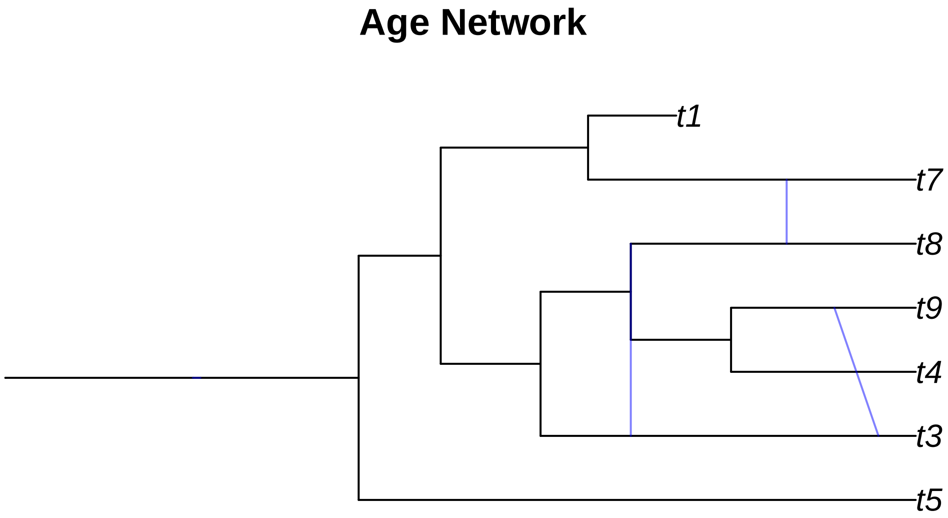


A few odd things happened here when we tried plotting the networks we simulated. We will go over these network by network.

- The **SSA network** only shows one hybridization event although we can see from the `reticulation` element that there are two events. This occurs when there is a hybridization where the two donor lineages have the same parental node.

- From the **GSA network** figure we can notice two issues. Firstly, it looks cut off or was plotting incorrectly. While the edge and reticulation matrix would confirm that the network is properly conformed, the plotting function of `ape` doesn't always render networks correctly. Secondly, the GSA network has a diagonal hybrid edge, and while this isn't improper, it does muddle the interpretation of the phylogeny. The diagonal line would seem to imply that there was gene flow from a lineage in the past to a more recent lineage. This clearly isn't the case as no lineages that we know of can time travel. These diagonal hybrid edges can result from incomplete lineage sampling, gene flow from a now extinct species, or lineage degenerative hybridization but it can still be hard to interpret diagonal edges.
- In the **Age network** we can see the same issue with one of the hybrid edges not appearing as in the **SSA network** and the diagonal edges as in the **GSA network**.

We correct these issues by using the `plottable.net()` function to modify the phylogeny to a more plotting-friendly network where gene flow always occurs between contemporary lineages. We aren't changing anything fundamental about the network, just how the edges get drawn.

```
ssa_pnet <-plottable.net(ssa_net)
gsa_pnet <-plottable.net(gsa_net)
age_pnet <-plottable.net(age_net)

plot(ssa_pnet,main="SSA Network")
plot(gsa_pnet,main="GSA Network")
plot(age_pnet,main="Age Network")
```

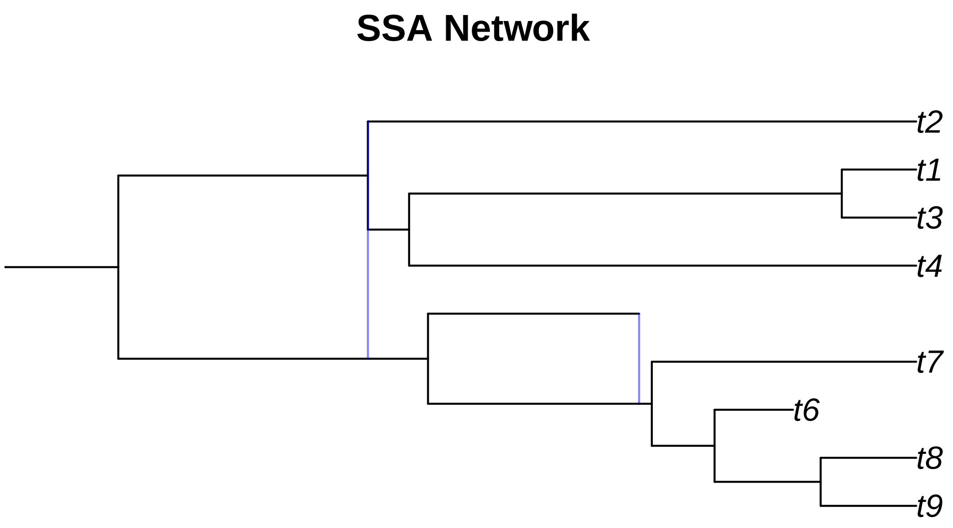

### GSA Network

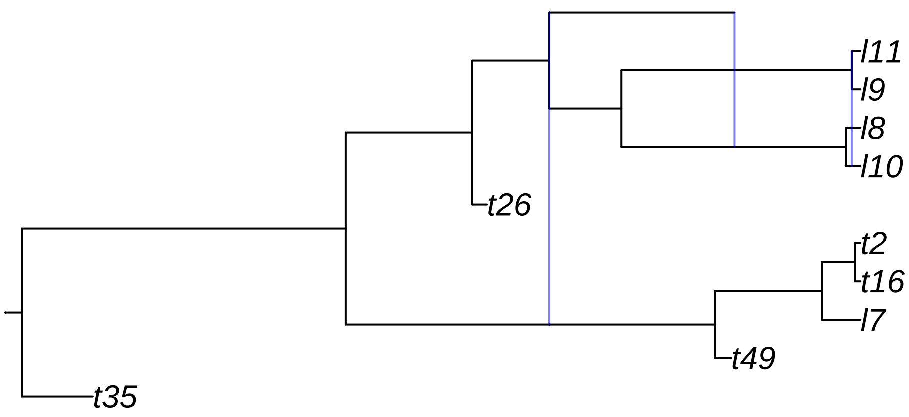


### Age Network

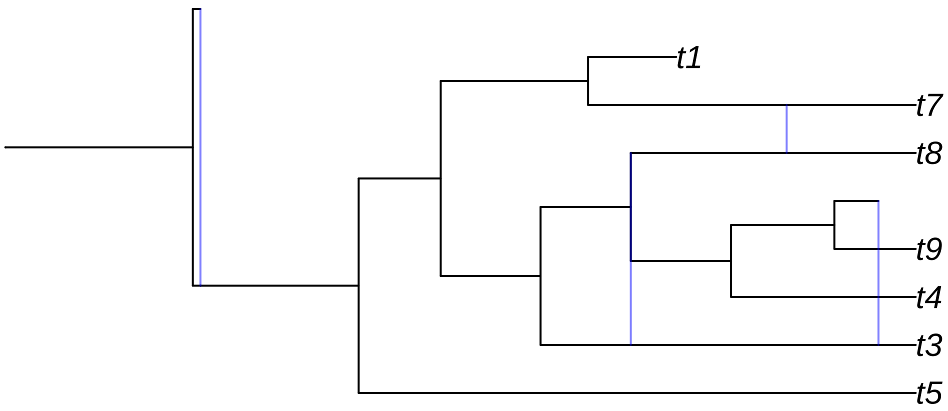


**Example 4 Network Classification**

Certain classes of phylogenetic networks have ideal properties. As such we may wish to know which classes our networks belong. Specifically, we can check whether our simulated networks are:

- **Tree-Child:** We can use `isTreeChild` to determine if a phylogenetic network is tree-child. A networks belongs to the tree-child class if for all internal nodes there exists at least one child node that is a tree-node *(Cardona, Pons, and Scornavacca 2019)*.
- **Tree-Based:** We can use `isTreeBased` to determine if a phylogenetic network is tree-based. A phylogenetic network is said to be tree-based if it can be constructed with a base tree that has additional linking arcs added *(Pons, Semple, and Steel 2019)*.
- **FU-Stable:** `isFUstable` can be used to determine if a phylogenetic network is FU-stable. A phylogenetic network is considered FU-stable if the unfolding and refolding of the network is isomorphic to the original network $N = F(U(N))$ *(Huber et al. 2016)*.
- **Level-k:** `getNetworkLevel` can be used to determine the level of the network. A level-k network has at least one biconnected component with k reticulations *(Gambette, Berry, and Paul 2009)*.

We can see these functions in action below:

```
isTreeChild(gsa_net)
#> [1] FALSE
isTreeBased(gsa_net)
#> [1] TRUE
isFUstable(gsa_net)
#> [1] FALSE
getNetworkLevel(gsa_net)
#> [1] 3
```

#### Example 5 Network Input And Output

Now that we've gotten a look at the networks we may wish to save these networks to file. We will be using the rich Newick Format for saving the phylogenies *(Cardona, Rosselló, and Valiente 2008; but see Wen et al. 2018)*. Notably, the `ape` function `write.evonet` does not save inheritance probabilities to file so we defined `write.net` for this purpose

```
write.evonet(ssa_net,file='') ##we can see that inheritance probabilities aren't included here
#> [1] "((((((((t9:0.08909672167,t8:0.08909672167):0.09893002096,t6:0.07316472041):0.05853474425,t7:0.2
my_newick<-write.net(ssa_net,file = '') ## if we include a file name the network will print to file ins
print(my_newick)
#> [1] "((((((((t9:0.08909672167,t8:0.08909672167):0.09893002096,t6:0.07316472041):0.05853474425,t7:0.2

my_net<-read.net(text=my_newick)
print(my_net)
#>
#>     Evolutionary network with 2 reticulations
#>
#>                  --- Base tree ---
#> Phylogenetic tree with 8 tips and 12 internal nodes.
#>
#> Tip labels:
#>   t9, t8, t6, t7, t2, t4, ...
#> Node labels:
#>   , , , , #H17, , ...
#>
#> Rooted; includes branch lengths.
str(my_net)##we can see that my_net has the inheritance element for inheritance probabilities
#> List of 7
```

```
#>  $ edge        : int [1:19, 1:2] 9 10 11 12 13 14 15 16 16 15 ...
#>  $ Nnode       : int 12
#>  $ tip.label   : chr [1:8] "t9" "t8" "t6" "t7" ...
#>  $ edge.length : num [1:19] 0.1053 0.2327 0.056 0.1968 0.0118 ...
#>  $ node.label  : chr [1:12] "" "" "" "" ...
#>  $ reticulation: int [1:2, 1:2] 12 18 13 17
#>  $ inheritance : num [1:2] 0.491 0.304
#>  - attr(*, "class")= chr [1:2] "evonet" "phylo"
```

### Reconstructed Phylogenies & Incomplete Sampling

Often not all extant tips on a phylogeny are sampled, nor do we have clear fossil records that tell us about the extinct species. We may wish to reflect this lack of knowledge by generating phylogenies with incomplete sampling or extinct lineages pruned. We can do this directly in the `sim.bdh` type functions by changing the `frac` and `stochsampling` arguments for incomplete sampling and by setting the `complete` argument to `FALSE` for the reconstructed phylogeny with extinct lineages removed. However, we can also do both of these actions post-hoc with the `incompleteSampling()` and `reconstructedNetwork()` functions.

#### Example 6: Post-hoc Incomplete Sampling an Reconstructed Networks

Our GSA network had a few extinct lineages, we can try pruning those with `reconstructedNetwork`. We will then randomly subsample 5 out of the 7 extant tips.

```
pruned_gsa <- reconstructedNetwork(gsa_net)
plot(plottable.net(pruned_gsa),main='Reconstructed Phylogeny')
pruned_gsa <- incompleteSampling(pruned_gsa,rho=5/7,stochastic = F)
plot(plottable.net(pruned_gsa),main='Reconstructed Phylogeny with Incomplete Sampling')
```

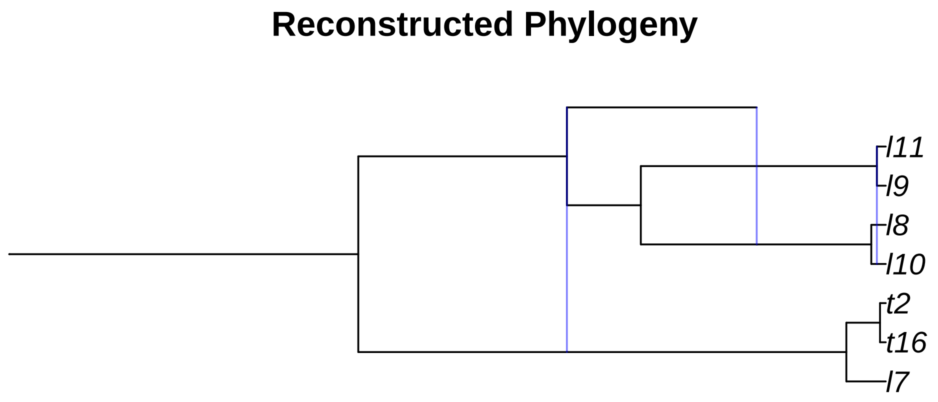

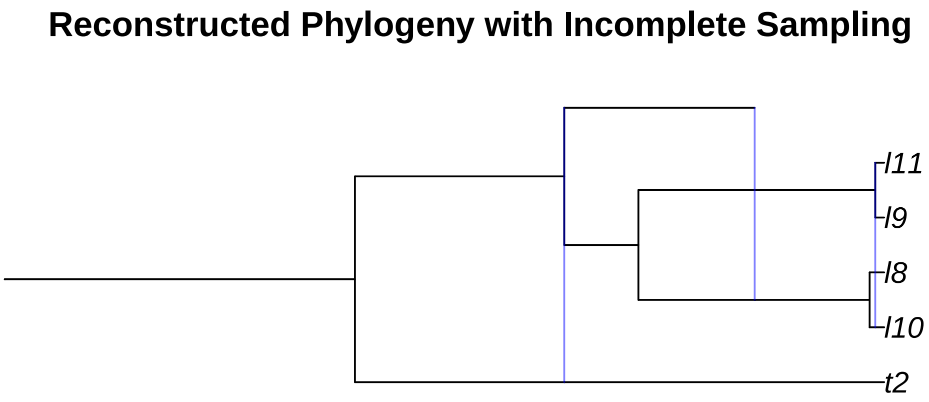


### More Complex Hybridization Models

We can extend our model of hybridization in two main ways:

1. **Make the probability of a successful hybridization a function of genetic distance.** We can do this to model the belief that it is less likely for more distantly related taxa to hybridize. This function will take in the genetic distance between two taxa and return the probability that the hybridization is successful. We use the same matrix of genetic distances that is defined in *Woodhams, Lockhart, and Holland (2016)*. Briefly, the genetic distance between two non hybrid taxa $X$ and $Y$ is defined as the summation of all edge lengths on the path from $X$ to $Y$. Genetic distances involving hybridized taxa is similar but hybridizations lead to multiple paths from $X$ to $Y$; in this case we use a weighted summation across all different paths where the weight for each path depends on the inheritance probabilities.
2. **Make successful hybridizations depend on some trait that also evolves along the tree.** This extension was made to model the belief that typically only lineages with the same ploidy and chromosome number can hybridize, although this can be generalized to model any discrete or continuous trait and have arbitrary rules for restricting hybridization.

#### Example 7A: Hybridizations Dependent On Genetic Distance

We can make the probability that a hybridization is successful by providing a function that is defined on the range $[0, \infty)$ for genetic distances and returns values from 0 to 1 for the probability of a successful hybridization. We have implemented several `make` functions for creating functions that fit these criteria. These functions are then used as the `hyb.rate.fxn` argument in the `sim.bdh` style functions.

```
##Here are some of the make functions
f1<-make.exp.decay(t=1,s=1)
f2<-make.linear.decay(threshold = 1)
f3<-make.stepwise(probs = c(1,0.5,0),distances = c(0.25,0.75,Inf))
f4<-make.polynomial.decay(threshold = 1,degree = 2)

##We can use any of these functions as the hyb.rate.fxn argument in a sim.bdh function
```

```
age_nets <-sim.bdh.age(age=2,numbsim=10,
                    lambda=1,mu=0.2,
                    nu=0.25, hybprops = hybrid_proportions,
                    hyb.inher.fxn = inheritance.fxn,
                    hyb.rate.fxn = f3)
```

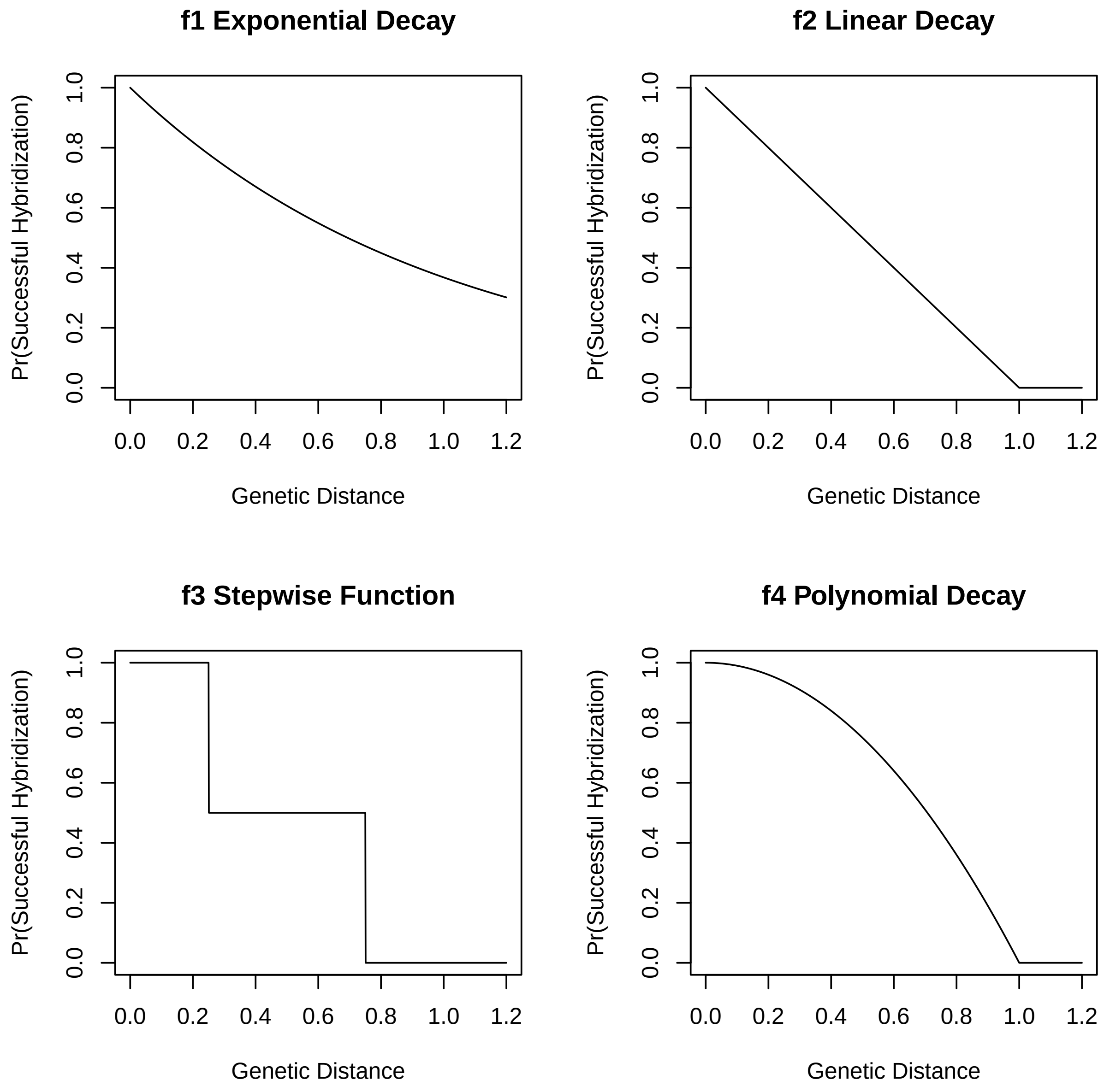


**Example 7B: Trait Dependent Hybridization**

A vast array of traits (both discrete and continuous) and mechanisms can control whether hybridization can occur between two species. We provide a general framework that gives the flexability to model many of these interactions. Later we give examples to showcase the some of the flavors of trait-dependent hybridization scenarios.

We can model trait dependent hybridization by supplying a list for the argument `trait.model` in the `sim.bdh` functions. This list contains initial value(s) for the trait at the beginning of the process, functions describing how the trait evolves, and a function that dictates whether taxa can hybridize based on the trait values between taxa. Specifically the list contains these named elements:

- **initial** The initial trait state on the phylogeny. **NOTE:** if the `MRCA` argument in the `sim.bdh` style functions is `TRUE`, 2 initial values will be needed since the process starts with 2 lineages.
- **hyb.event.fxn** A function that denotes the trait of a hybrid child after a hybridization event. The function should have the arguments `parent_states` and `inheritance`. `parent_states` is vector with the ploidy states of the hybrid parents while `inheritance` is the inheritance probability of the first lineage denoted in `parent_states`.
- **hyb.compatibility.fxn** A function that describes when hybridization events can occur between two taxa based on their traits. The function should have the argument `parent_states`, a vector with the trait states of the two parents to the hybrid child. The function should return `TRUE` for when a hybridization event is allowed to proceed and `FALSE` otherwise.
- **time.fxn** A function that describes how traits change over time. The function should have the arguments `poly_states` and `timestep` in that order. `poly_states` is a vector containing the ploidy of all taxa while `timestep` is the amount of time given for trait evolution. The function should return a vector with the updated ploidy states of all taxa.
- **spec.fxn** A function that describes how the trait changes at speciation events.The function should have the argument `tip_state` which has the state of the lineage just before speciation. The function should return a vector with two values, one denoting the trait of each of the two new species after the event.

Here we give an example of simple ploidy evolution that considers autopolyploidy and allopolyploid hybridization. We restrict allopolyploidy events to only occur between lineages with the same ploidy.

```
initial_val<-2 ## The root starts off at 2N

###function for what happens at hybridization event
hyb_e_fxn <- function(parent_states,inheritance){
  ##For allopolyploidy we add the ploidy of both parents
  return(sum(parent_states))
}

##Function for determining whether hybridization occurs
hyb_c_fxn <-function(parent_states,hybrid_state){
  ##Hybridization occurs only when the ploidy is the same
  return(parent_states[1]==parent_states[2])
}


##Function for how the trait changes over time
t_fxn <- function(trait_states,timestep){
  ##We assume that autopolyploidy occur exponentially with rate lambda
  lambda<- 2 ##Rate of autopolyploidy

  ##The number of autopolyploidy events that occur on each lineage over the timestep
  nevents<-rpois(length(trait_states),timestep)

  ##each event doubles the ploidy
  new_states<- trait_states * (2^nevents)
```

```
  return(new_states)
}

##Function for how the trait changes at speciation events
s_fxn <-function(tip_state){
  ##Ploidy doesn't change at speciation events.
  ##Both daughter lineages have the same ploidy as the parent
  return(c(tip_state,tip_state))
}

trait_model<-make.trait.model(initial_states = initial_val,
                              hyb.event.fxn = hyb_e_fxn,
                              hyb.compatibility.fxn = hyb_c_fxn,
                              time.fxn = t_fxn,
                              spec.fxn = s_fxn)


trait_nets <-sim.bdh.age(age=2,numbsim=10,
                    lambda=1,mu=0.2,
                    nu=0.25, hybprops = hybrid_proportions,
                    hyb.inher.fxn = inheritance.fxn,
                    trait.model = trait_model)
```

Next we give an example of a continuous trait evolution where the trait dissimilarity determines hybrid compatibility between two species. This might reflect something like differences in body size mechanically isolating the two species or the time of the mating seasons creating temporal isolation. We choose to take a probabilistic approach by saying, much like the genetic-distance dependent hybridization, that there is some decay function that relates hybridization success to trait dissimilarity.

```
initial_val<-0 ## The root starts off at 0

###function for what happens at hybridization event
hyb_e_fxn <- function(parent_states,inheritance){
  ## Take a weighted average of the two traits
  return(sum(parent_states*c(inheritance,1-inheritance)))
}

##Function for determining whether hybridization occurs
hyb_c_fxn <-function(parent_states,hybrid_state){
  #Make linear decay function that decreases hybrid success probability linearly
  #Hybridization cannot occur when the traits differ by more than 4
  decay.fxn <- make.linear.decay(4)

  #Trait dissimilarity
  diss<-abs(parent_states[1]-parent_states[2])

  success_prob<-decay.fxn(diss)
  return(runif(1,0,1)<=success_prob)
}


##Function for how the trait changes over time
t_fxn <- function(trait_states,timestep){
```

```
  ##We assume brownian motion on the continuous trait
  sigma<- 2 ##sqrt(Rate of evolution)

  ##Make brownian motion draws for each lineage
  delta_x <- rnorm(length(trait_states),mean=0,sd=sigma*sqrt(timestep))

  return(delta_x+trait_states)
}

##Function for how the trait changes at speciation events
s_fxn <-function(tip_state){
  ##No change at speciation
  return(c(tip_state,tip_state))
}

trait_model<-make.trait.model(initial_states = initial_val,
                              hyb.event.fxn = hyb_e_fxn,
                              hyb.compatibility.fxn = hyb_c_fxn,
                              time.fxn = t_fxn,
                              spec.fxn = s_fxn)


trait_nets <-sim.bdh.age(age=2,numbsim=10,
                    lambda=1,mu=0.2,
                    nu=0.25, hybprops = hybrid_proportions,
                    hyb.inher.fxn = inheritance.fxn,
                    trait.model = trait_model)
```

Lastly, we model a case where hybridization can only occur if the hybrid species is sufficiently different enough from the parental ones. This might be used to model hybrid breakdown and the hybrid lineage needing to exploit a different niche than the parental lineages due to a lower fitness. We also decide to allow transgressive trait evolution in the hybrid lineages. The trait in the hybrid lineage is not always an intermediate value between the two lineages but sometimes something else entirely, we model that here.

```
initial_val<-0 ## The root starts off at 0

###function for what happens at hybridization event
hyb_e_fxn <- function(parent_states,inheritance){
  ## Take a weighted average of the two traits
  hyb_val<-sum(parent_states*c(inheritance,1-inheritance))

  ##Now allow transgressive evolution by making a random draw from a normal distribution
  transgression <- rnorm(1,0,4)

  return(hyb_val+transgression)
}

##Function for determining whether hybridization occurs
hyb_c_fxn <-function(parent_states,hybrid_state){
  ##Lets say that anything outside 10% of the hybrid lineage trait value is a different nich
  niche_bound <-hybrid_state*0.1

  lower_bound <-hybrid_state-niche_bound
```

```r
  upper_bound <-hybrid_state+niche_bound

  ##Check if the parental lienages are within the niche boundary
  if(all( (parent_states < lower_bound) | (parent_states > upper_bound) )){
    ##Parent lineages are outside the hybrid niche. Hybrid survives
    return(TRUE)
  }else{
    ##Parent lineage inside the hybrid niche. Hybrid breakdown
    return(FALSE)
  }

}


##Function for how the trait changes over time
t_fxn <- function(trait_states,timestep){
  ##We assume brownian motion on the continuous trait
  sigma<- 2 ##sqrt(Rate of evolution)

  ##Make brownian motion draws for each lineage
  delta_x <- rnorm(length(trait_states),mean=0,sd=sigma*sqrt(timestep))

  return(delta_x+trait_states)
}

##Function for how the trait changes at speciation events
s_fxn <-function(tip_state){
  ##No change at speciation
  return(c(tip_state,tip_state))
}

trait_model<-make.trait.model(initial_states = initial_val,
                              hyb.event.fxn = hyb_e_fxn,
                              hyb.compatibility.fxn = hyb_c_fxn,
                              time.fxn = t_fxn,
                              spec.fxn = s_fxn)


trait_nets <-sim.bdh.age(age=2,numbsim=10,
                    lambda=1,mu=0.2,
                    nu=0.25, hybprops = hybrid_proportions,
                    hyb.inher.fxn = inheritance.fxn,
                    trait.model = trait_model)
```

pcbi.1007347.

Cardona, Gabriel, Francesc Rosselló, and Gabriel Valiente. 2008. "Extended Newick: It Is Time for a Standard Representation of Phylogenetic Networks." *BMC Bioinformatics* 9 (1): 532. https://doi.org/10.1186/1471-2105-9-532.

Davín, Adrián A., Théo Tricou, Eric Tannier, Damien M. de Vienne, and Gergely J. Szöllosi. 2020. "Zombi: A Phylogenetic Simulator of Trees, Genomes and Sequences That Accounts for Dead Linages." *Bioinformatics* 36 (4): 1286–88. https://doi.org/10.1093/bioinformatics/btz710.

Gambette, Philippe, Vincent Berry, and Christophe Paul. 2009. "The Structure of Level-k Phylogenetic Networks." In *Annual Symposium on Combinatorial Pattern Matching*, 289–300. Springer.

Hagen, Oskar, and Tanja Stadler. 2018. "TreeSimGM: Simulating Phylogenetic Trees Under General Bellman–Harris Models with Lineage-Specific Shifts of Speciation and Extinction in r." *Methods in Ecology and Evolution* 9 (3): 754–60. https://doi.org/https://doi.org/10.1111/2041-210X.12917.

Hartmann, Klaas, Dennis Wong, and Tanja Stadler. 2010. "Sampling Trees from Evolutionary Models." *Systematic Biology* 59 (4): 465–76. https://doi.org/10.1093/sysbio/syq026.

Höhna, Sebastian. 2013. "Fast simulation of reconstructed phylogenies under global time-dependent birth–death processes." *Bioinformatics* 29 (11): 1367–74. https://doi.org/10.1093/bioinformatics/btt153.

Höhna, Sebastian, Michael R. May, and Brian R. Moore. 2015. "TESS: an R package for efficiently simulating phylogenetic trees and performing Bayesian inference of lineage diversification rates." *Bioinformatics* 32 (5): 789–91. https://doi.org/10.1093/bioinformatics/btv651.

Huber, Katharina T., Vincent Moulton, Mike Steel, and Taoyang Wu. 2016. "Folding and Unfolding Phylogenetic Trees and Networks." *Journal of Mathematical Biology* 73 (6): 1761–80.

Mallet, James, Nora Besansky, and Matthew W. Hahn. 2016. "How Reticulated Are Species?" *BioEssays* 38 (2): 140–49. https://doi.org/10.1002/bies.201500149.

Morin, M. M., and B. M. E. Moret. 2006. "NetGen: Generating Phylogenetic Networks with Diploid Hybrids." *Bioinformatics* 22 (15): 1921–23. https://doi.org/10.1093/bioinformatics/btl191.

Nee, Sean. 2006. "Birth-Death Models in Macroevolution." *Annual Review of Ecology, Evolution, and Systematics* 37 (1): 1–17. https://doi.org/10.1146/annurev.ecolsys.37.091305.110035.

Pons, Joan Carles, Charles Semple, and Mike Steel. 2019. "Tree-Based Networks: Characterisations, Metrics, and Support Trees." *Journal of Mathematical Biology* 78 (4): 899–918.

Stadler, Tanja. 2011. "Simulating Trees with a Fixed Number of Extant Species." *Systematic Biology* 60 (5): 676–84. https://doi.org/10.1093/sysbio/syr029.

Wen, Dingqiao, Yun Yu, Jiafan Zhu, and Luay Nakhleh. 2018. "Inferring Phylogenetic Networks Using PhyloNet." *Systematic Biology* 67 (4): 735–40. https://doi.org/10.1093/sysbio/syy015.

Woodhams, Michael D., Peter J. Lockhart, and Barbara R. Holland. 2016. "Simulating and Summarizing Sources of Gene Tree Incongruence." *Genome Biology and Evolution* 8 (5): 1299–1315. https://doi.org/10.1093/gbe/evw065.

Zhang, Chi, Huw A. Ogilvie, Alexei J. Drummond, and Tanja Stadler. 2018. "Bayesian Inference of Species Networks from Multilocus Sequence Data." *Molecular Biology and Evolution* 35 (2): 504–17. https://doi.org/10.1093/molbev/msx307.
